# Androgen-mediated TGFβ expression suppresses anti-tumor neutrophil response in bone metastatic prostate cancer

**DOI:** 10.1101/2022.12.30.522329

**Authors:** Massar Alsamraae, Diane Costanzo-Garvey, Benjamin A. Teply, Shawna Boyle, Gary Sommerville, Zach Herbert, Colm Morrissey, Alicia J. Dafferner, Maher Y. Abdalla, Rachel W. Fallet, Tammy Kielian, Heather Jensen-Smith, Edson I. deOliveira, Keqiang Chen, Ian A. Bettencourt, Ji Ming Wang, Daniel W. McVicar, Tyler Keeley, Fang Yu, Leah M. Cook

**Author notes:** **Corresponding Author**: Leah M. Cook, Dept. of Pathology/Microbiology, University of Nebraska Medical Center, 985900 Nebraska Med Center, Omaha, NE, USA 68198. Designates equal contribution (Massar Alsamraae, Diane Costanzo-Garvey).

## Abstract

Prostate Cancer (PCa) bone metastases are associated with spinal cord compression, fracture, bone pain and death. Despite advances in the medical therapy for localized disease, metastatic disease is incurable and osseous progression is largely dictated by tumor-stromal interactions in the bone microenvironment. We showed previously that tumor bone neutrophils are tumoricidal to PCa but lose their cytotoxic potential as the tumor progresses. However, there have been no studies to date to clinically define and characterize neutrophils throughout the prostate cancer disease spectrum to determine their biomarker potential. Using patient peripheral blood polymorphonuclear neutrophils (PMNs), we identify that PCa progression dictates PMN properties, including viability, cell surface markers and gene expression. However, the majority of PMNs elicited an anti-tumor response *ex vivo* demonstrating that PMN cytotoxicity is cell autonomous and independent of PCa disease stage. In fact, we identify a novel role for androgen regulation, i.e., androgen deprivation therapy (ADT), in suppressing PMN cytotoxicity via altered transforming growth factor beta receptor I (TβRI). Using preclinical models, we found that high dose testosterone/bipolar androgen therapy (BAT) and genetic or pharmacologic TβRI inhibition combined with standard ADT rescued the androgen-associated suppression and restored PMN anti-tumor immune response. This combination provided a therapeutic strategy more impactful than ADT alone, in bone metastatic prostate cancer (BM-PCa). These studies: 1) highlight a necessity for both molecular and functional characterization of PMNs per cancer type and 2) reveals the ability to program PMN immune response for successful targeting of BM-PCa.

## Introduction

Prostate cancer (PCa) is the second most common cause of cancer death among men in the United States, accounting for an estimated 34,000 cancer deaths in 2022, which are mostly associated with metastatic disease^1,2^. The current standard of therapy for advanced prostate cancer is either prostatectomy or initial androgen deprivation therapy (ADT) with response to therapy being initially characterized by biochemical response e.g., changes in the levels of prostate-specific antigen (PSA)^3,4^. Approximately 50% of PCa patients progress to advanced stage disease characterized by biochemical recurrence and often, progression to metastatic disease or metastatic castration-resistant prostate cancer (mCRPC)^5^. Although mCRPC patients exhibit progressing disease, many will respond to additional androgen receptor (AR) directed therapy given in the form of more selective androgen inhibitors including, androgen synthesis inhibitors and androgen receptor (AR) inhibitors, such as abiraterone acetate (which inhibits CYP17A1) and enzalutamide, respectively^6–8^. Even with a ~4-6 month improvement in overall survival, mCRPC patients typically succumb to the disease within 3 years after diagnosis of metastases. These statistics demonstrate a significant need to identify additional therapies or personalized therapeutic approaches.

Bone is the most frequent tissue site for metastatic prostate cancer (mPCa). It has been classically shown that within the bone microenvironment mPCa promotes bone remodeling, e.g., excessive bone degradation, which subsequently results in the release of bone-sequestered growth factors such as transforming growth factor beta, TGFβ, which further promotes tumor growth and a continuous cycle of PCa growth in bone^9,10^. Importantly, PCa interactions with bone stromal cells contribute significantly to the progression of mPCa, however it is still unclear why bone-targeted therapies, such as anti-RANKL inhibitors and bisphosphonates have little to impact on patient survival, respectfully^11,12^. In addition to bone cells, including bone degrading osteoclasts and bone forming osteoblasts, which surprisingly comprise only ~1% of cells within the bone compartment, there is emerging evidence that other cells in the bone marrow can regulate PCa progression^13^. Notably, we previously showed that bone marrow neutrophils and neutrophil precursors, which account for ~50-60% of cells in bone marrow^13,14^, migrate to BM-PCa and are protective against growth of mPCa in bone.

Neutrophils are the most abundant of leukocytes population, representing 50–70% of all immune cells. Humans produce more than 10^11^ neutrophils daily. In bone marrow, neutrophils are generated and released into circulation daily, ready to elicit an innate immune response to pathogens and tissue damage. Thus, a large storage pool of neutrophils consistently remains in the bone marrow. There are conflicting data of the role of neutrophils in tumor progression, demonstrating both anti-tumoral/ “N1-like” and or protumor/ “N2-like” functions that appear to be dictated by the specific tissue milieu^15^. However, the role of neutrophils in the progression of bone metastatic prostate cancer (BM-PCa) is still unclear.

We previously found that bone marrow neutrophils target prostate cancer growth through inhibition of tumor cell STAT5 expression; however, PCa tumors in the bone quickly acquired resistance to neutrophils and neutrophil cytotoxicity was diminished, though not switched to a pro-tumoral role^16^. Understanding the mechanisms that drive anti-tumor neutrophil immune response can be utilized to develop novel targets for treating BM-PCa. The goal of this study was to comprehensively define the effects of PCa on neutrophil function throughout progression. To do this, we utilized patient-derived peripheral blood neutrophils for comparison to primary bone marrow neutrophils and ran a preclinical trial which revealed several novel insights into neutrophils in PCa. Importantly, we observed classical behaviors of tumor-associated neutrophils (TANs), novel characteristics of tumor-bone neutrophils (TBNs), and also a therapy-associated regulation of neutrophil cytotoxicity that was independent of disease stage. Thus, we reveal a novel mechanism through which androgens regulate neutrophil function and identify PCa-associated neutrophil properties that can be utilized for development of neutrophil-focused cancer immunotherapy.

## Materials & Methods

### Patient sample collection

Healthy men and PCa patients were deemed eligible and consented for sample donation within the UNMC integrated Cancer Repository for Cancer Research (iCaRe2). The iCaRe2 is a multi-institutional resource created and maintained by the Fred & Pamela Buffett Cancer Center to collect and manage standardized, multi-dimensional, longitudinal data and biospecimens on consented adult cancer patients, high-risk individuals, and normal controls. Patients for this study were specifically recruited from the Genitourinary Cancer Registry (GU-CARE), and all samples collected under the guidance and approval of the UNMC Institutional Review Board (IRB) under iCaRe2. The study consisted of 4 groups (n= >17/group): 1) healthy (no cancer or pathological disease) (n=17), 2) localized PCa (n=22), 3) bone metastatic hormone-/castration-sensitive PCa (n=18), and 4) bone metastatic castration-resistant PCa (n=20). Patient blood was collected and PMNs immediately isolated after collection and utilized for downstream analyses, including: cell counts, viability, flow cytometry, RNA collection, RNA sequencing, and co-culture assay. Although some patients presented with visceral metastases, the co-culture data in this manuscript includes only bone metastatic patients. PCa patients received differing therapies, including: Leuprolide, hormone signaling inhibitors (ARSIs; abiraterone, enzalutamide, darolutamide, apalutamide) and chemotherapy (docetaxel). Other bone-targeted therapies included radium-223,zoledronic acid and denosumab. All rapid autopsy metastatic tumors were from patients who signed written informed consent under the aegis of the Prostate Cancer Donor Program at the University of Washington (IRB protocol # 2341).

### Mice

Only male mice were used for these studies. For neutrophil isolations, C57BL/6 mice were utilized (Jackson Laboratory; # 000664). Mice (8-12 weeks) were used for all experiments using primary neutrophils and mouse RM1 *in vivo* intratibial metastasis models. For human PCa cell intratibial metastasis models, immunocompetent SCID/Beige (C.B-17/IcrHsd-PrkdcscidLystbg-J) mice were utilized (Envigo). *TβRI^flox/flox^* mice were purchased (Jackson Laboratory; #028701) and backcrossed 8 times to C57BL/6 to obtain a pure background. Neutrophil-selective *TβRI* knockout mice, Catchup mice^17^ (Ly6G^*Cre/dtom*^) were crossed with *TβRI^flox/flox^* mice; knockout *TβRI^−/−^* was verified using immunoblot analysis of mouse tissues. Mice were housed on a 12 hour light/dark cycle, with free access to food and water. All procedures performed were approved by IACUC (UNMC).

### Cell culture media, reagents, and buffers

Neutrophil isolation buffer consists of 1X PBS, 2% FBS and 2 mM EDTA. RPMI complete media consists of RPMI (Hyclone), 10% FBS (Peak Serum), and 1% penicillin/ streptomycin. DMEM complete media consists of DMEM (Hyclone), 10% FBS (Peak Serum), and 1% penicillin/streptomycin. Luciferase-expressing C42B and PAIII were supplemented with 10 μg of puromycin, and LNCaP with 5 μg puromycin (Gibco). For TβRI assays, primary neutrophils were incubated in CM supplemented with RepSox ((5 nM; Selleckchem). To collect conditioned media (CM), cell lines were washed with Phosphate Buffered Saline (PBS) to remove serum, full medium was replaced with serum-free medium RPMI and cells were incubated for 24 hours. CM was collected and centrifugation was done to remove cellular debris and stored at 4 °C until usage. Total protein content of CM was measured using BCA assay (ThermoFisher) to ensure equal protein concentrations for treating neutrophils. Fresh media was collected every 2 weeks for experimental use.

### Human neutrophil isolation

From patients, peripheral blood neutrophils were isolated from freshly collected blood, within 5 hours of collection. Neutrophils were isolated using a commercially available kit (EasySep Human Neutrophil Isolation; StemCell). For mouse tumor-associated neutrophils (TANs) and primary neutrophil co-cultures, bone marrow neutrophils were isolated from mouse tibia and femurs of male C57BL/6 mice. Bones were cleared of tissue and muscle, and the epiphysis was removed and discarded. A hole was made in the bottom of a 0.65 mL tube, and one bone was placed individually per tube. This tube was placed into a 1.5-mL tube for bone marrow collection, where it was then centrifuged at high speed for < 5 s. The bone marrow was re-suspended in 1 mL of neutrophil isolation buffer and filtered using a 70 micron filter. Bones from each mouse were pooled and counted, and the MojoSort Neutrophil Enrichment (Biolegend) protocol was followed, as per manufacturers’ instructions. All neutrophil described functional assays utilized mouse bone marrow neutrophils.

### Real-time Glo MT cell viability assay

Neutrophils were incubated in triplicate at 100,000 per well in PCa CM with 2x Real-Time Glo reagent (Promega). MT Cell Viability Substrate and NanoLuc® Enzyme were added in equal volumes to culture media to create the 2x Real-time Glo reagent. This media was added directly to the cells at time zero, and luminescence was read at the indicated time points using a luminometer. Mouse neutrophils were treated for 2 hours with CM with enzalutamide at 3 μM and 10 μM, then the luciferase was quantified at 3, 6, 18 and 24 hours.

### Flow cytometry

From patient samples, isolated peripheral blood neutrophils were washed and reconstituted in FACS buffer (2% FBS in 1X PBS) at 1×10^6^ cells in 200 μL. For tumor studies, bone marrow was flushed from tumor-bearing and saline-injected tibiae using a syringe and excess cells were further flushed out of the marrow with FACS buffer. For staining, cells were incubated on ice with fluorophore-conjugated antibodies added at a maximum of 1 μL per 10^6^ cells in appropriate antibodies listed in Supplemental Table 1. Cell viability dye, Live/Dead (Invitrogen), was also added at a concentration of 0.2 μL per 10^6^ cells. Stained cells were incubated with antibody on ice in the dark for 20 min, rinsed with 1X PBS and were fixed in 1% formaldehyde in 1XPBS for 30 min in the dark. Prior to analysis, cells were reconstituted in FACS buffer. For all analyses, single and live cells were gated and percentage of positive cells per marker was quantified.

### RNA Library Preparation and Sequencing

For patient samples, peripheral blood neutrophils were isolated and reconstituted in Trizol reagent. RNA was isolated from Trizol using DirectZol kit, according to manufacturer instructions. Libraries were prepared using Roche Kapa Biosystems RiboErase and RNA HyperPrep sample preparation kits from 100ng of RNA. RNA samples were fragmented at 94C for 8 minutes with 14 cycles of PCR post-adapter ligation according to manufacturer’s recommendation. The finished dsDNA libraries were quantified by Qubit fluorometer and Agilent TapeStation 2200. Libraries were pooled in equimolar ratios and evaluated for cluster efficiency and pool balance with shallow sequencing on an Illumina MiSeq. Final sequencing was performed on an Illumina NovaSeq with paired-end 100bp reads at the Dana-Farber Cancer Institute Molecular Biology Core Facilities. *RNASeq Analysis*.Sequenced reads were aligned to the UCSC hg38 reference genome assembly and gene counts were quantified using STAR (v2.7.3a)^18^. Differential gene expression testing was performed by DESeq2 (v1.22.1)^19^. Initial RNAseq analysis was performed using the VIPER snakemake pipeline^20^. Data was further analyzed using Ingenuity Pathway Analysis software (IPA; Qiagen). The full gene list was uploaded into the NCBI Repository and can be accessed at GEO Accession: GSE197609.

### Real-time qPCR

For investigating the effects of enzalutamide, mouse neutrophils were incubated for 30 minutes at 37 °C with enzalutamide (3 μM and 10 μM) compared to vehicle control (DMSO), and gene expression was performed using Bio-Rad CFX Real-Time System. For analysis of gene expression, Trizol reagent was added to treated neutrophils and RNA was extracted using the standard Trizol isolation protocol. RNA (1 μg) was used to synthesize cDNA using qSCRIPT Super mix (Quantabio) and PCR was performed using Perfecta SYBR Green FastMix (Quantabio). PCR was run using Bio-Rad CFX Real-Time System. PCR conditions were as follows for all primer sequences (Integrated DNA Technologies; sequences listed in **Supplemental Table 2**): Step 1: 95° 30s; Step 2: 95° 5s, 57° 15s, 72° 10s, 95° 10s (×39 cycles); Step 3: Melt curve 65° to 95° at increments of 0.5° for 5s. For enzalutamide experiments: mouse primary bone marrow neutrophils were treated with 3 μM and 10 μM for 2 hours RNA was isolated from neutrophils which had been incubated for 3 hours at 37 °C, in PCa CM supplemented with 2% FBS.

### PCa:Neutrophil Co-culture assay

PCa co-cultures with neutrophils Luciferase-expressing LNCaP or C42B cells were plated at 40,000 cells/well in a 24-well plate, in triplicate per condition. Twenty-four hours later, neutrophils were isolated, re-suspended in complete medium, and plated in direct contact with cancer cells at a ratio of 10:1 (neutrophils/cancer) and 3 μM, 10 μM enzalutamide were added into the media or neutrophils were treated with enzalutamide for 30 minutes prior to addition to the culture where noted. After incubation overnight, neutrophils were removed, and cancer cell viability was measured with Trypan Blue Exclusion assay, using a hemacytometer.

### Neutrophil Extracellular Trap (NET) Production assay

For analysis of PMN NET secretion, primary mouse neutrophils were incubated for 2 hours in prostate CM (LNCaP, C42B) with or without Enzalutamide/ Abiraterone, or complete RPMI supplemented with PMA (100 nM) as a positive control. Sytox Green dye (500 nM; Sigma) was added to each condition, and after 30 min, images were taken of each well by an EVOS FL Auto microscope. The number of green fluorescent DNA traps/NETs was measured as a percentage of total cells to distinguish between dying cells that absorbed the Sytox Green dye. CM-treated neutrophils were compared to neutrophils incubated in serum-free media RPMI.

### Neutrophil migration assay

Primary neutrophils were isolated from mouse hind limbs using MojoSort and 1×10^5^ seeded in the top of transwell migration chambers, with a pore size of 5 microns (Costar; Ref # 3422). Neutrophils were allowed to migrate towards prostate cancer CM added to the bottom of the respective wells for one hour. To examine impact of androgen signaling, mouse neutrophils were pre-treated with 3 μM of Enzalutamide or Abiraterone for 30 minutes before addition the insert. Neutrophils were allowed to migrate towards specific conditions, for 1 hour: serum-free media, serum containing 2% FBS, serum-free LNCaP conditioned media (CM), and serum-free C42B CM. Inserts were fixed in 100% methanol for 5 minutes, washed in PBS to remove non-adherent cells and stained with hematoxylin for quantitation of the number of migrating neutrophils per insert. Insert membranes were mounted on a slide using Permount (Fisher Scientific) and imaged using an EVOS FL Auto microscope (Invitrogen; AMAFD1000) at 10x magnification.

### Immunoblot analysis

Protein was isolated from patient PMNs using RIPA buffer (Thermofisher, MA) and total protein quantified using BCA assay (Pierce, Thermofisher, MA). Protein (50 μg) was run on a 12% SDS gel and then transferred to a PVDF membrane. The membrane was stained with 0.1% (w/v) Ponceau S in 5% (w/v) acetic acid for 5 minutes at room temperature. Background staining was removed with 3 washes of distilled water prior to capturing an image. Distilled water was used to destain the membrane, non-specific binding was blocked with 5% non-fat milk in TBST for 1 hour at room temperature followed by overnight incubation with the primary antibody, 1:1000 in 5% non-fat milk, at 4°C. The following primary antibodies were used: MnSOD (#06-984, MilliporeSigma, Burlington MA), Catalase (#ab76024, Abcam, Boston MA), CuZnSOD/SOD1 (#ab51254, Abcam, Boston MA), GAPDH (#sc-32235, Santa Cruz Biotechnology, Dallas TX). The blot was washed in TBST prior to incubating with the secondary antibody, 1:4700 in 5% non-fat milk (anti-rabbit IgG #ADI-SAB-300-J, Enzo Lifesciences, Farmingdale NY or anti-mouse IgG #7076, Cell Signaling Technology, Danvers MA). The membranes were imaged using Azure Biosystems Radiance Plus substrate solution and an Azure c600 Imaging System.

### Immunohistochemistry

Patient bone specimens and mouse tibia bone sections were dewaxed and hydrated through an alcohol gradient. Antigen retrieval for human and mouse specimens was performed using Tris-EDTA buffer (pH 9) in a pressure cooker on high temp for 6 minutes. Tissues were then blocked in 10% serum in 1X tris-buffered saline (TBS) for 1 hour prior to overnight incubation in primary antibodies (Myeloperoxidase (7.5 μg/ml), R&D MAB3174; TGFβ Receptor I (1:100), Millipore, ABF17-1). Following washes, species-specific secondary AlexaFluor antibodies were incubated 1:1000 on the tissues for 1 hour at room temperature. Fluorescent images were taken on a Zeiss Axio Imager Z2 at 20x.

### In vivo mouse models

#### Androgen regulation study

To examine the impact of androgen regulation on tumor burden and TAN function, we performed a preclinical trial using standard-of-care therapy for BM-PCa, enzalutamide, and bipolar androgen therapy (BAT). Male SCID Beige mice were treated by subcutaneous injection once with Degarelix (commercially available gonadotropin releasing hormone antagonist; 10mg/kg) as initial ADT. The following day, mice were injected intratibially with C42B prostate cancer cells. Subcutaneous pellet placement will be performed 1 week later (to allow for confirmed tumor take). For injections, luciferase-expressing C42B, were grown to confluence, trypsinized and washed with 1X PBS, and filtered using a 70-μM nylon filter. Cells were counted and reconstituted for intratibial injection of 5×10^5^ C42B per 40 μL volume per mouse. Mice were anesthetized with isoflurane, and 5×10^5^ C42B injected into the right tibia. An equal volume of PBS was injected into the contralateral limb, as a control for the intratibial injection. At day 3 post-intratibial injection, tumor burden was imaged via bioluminescence. To do this, mice were given 10 μL/g of D-luciferin (15 mg/mL) (Gold Bio) intraperitoneally and imaged using the IVIS Spectrum imager (PerkinElmer). Luciferase signal was quantified 15 min after injection, using the Living Image Software per manufacturer’s instructions. Using bioluminescence intensity, mice were randomized into 6 treatment groups (n = 4-5/group; based on available mice after confirmed tumor take) to receive either: 1) enzalutamide (10mg/kg) or 2) vehicle control; 3) placebo subcutaneous pellet, and 4) slow-release 5-α-dihydrotestosterone pellet (DHT;12.5mg 21-day release; Innovative Research); 5) enzalutamide plus placebo pellet and 6) enzalutamide plus DHT pellet. Pellets were subcutaneously transplanted 7 days after the start of Enzalutamide treatment to allow for confirmed tumor take. IVIS imaging was used for longitudinal measurement of tumor burden. Mice were euthanized at 2 weeks to allow for isolation of bone marrow tumor-bone neutrophils (TBNs). Tumor-bearing and saline-injected tibia were flushed with sterile PBS and TBNs isolated using MojoSort Neutrophil Isolation kit and for analysis in co-culture and for protein analysis. Protein was isolated from TBNs using RIPA Lysis buffer, total protein content measured using BCA assay, and immune-modulating cytokines and chemokines examined by protein array (RayBiotech; Mouse Cytokine Array C3), according to protocol. As a control, tumor naïve neutrophils (from tibia injected with saline) were used for comparison. Flow cytometry was performed on neutrophils from tumor bearing and tumor naïve mice using the following markers: CD11b, CD45, Ly6C, Ly6G, F4/80 and CD11c (Biolegend).

#### TGFβ knockout and inhibitor study

To examine the impact of neutrophil-selective knockout of TGFβ receptor I on BM-PCa *in vivo*, the mouse RM1 prostate cancer model was utilized; 3.5 x 10^4^ luciferase-expressing RM1 were injected into tibia of 6-week old TβRI knockout (*TβRI^−/−^*) or control wildtype TβRI mice (*TβRI^flox/flox^*). Additionally, RM1 was used to examine the outcome of dual (and single) TβRI and AR inhibition on PCa growth in bone. For this study, 3.5 x 10^5^ luciferase-expressing RM1 were injected into tibia; 3 days post-injection, mice were randomized via bioluminescence into 4 treatment groups (n=5-7/group based upon confirmation of tumor take): 1) vehicle control (DMSO), 2) enzalutamide (10mg/kg), 3) RepSox, TβRI inhibitor (5mg/kg), and 4) combination Enzalutamide and TβRI inhibitor. For both studies, tumor burden was measured longitudinally throughout the study using bioluminescence and hind limbs collected at the end of the study for downstream analyses.

### Two-photon microscopy

An upright Olympus FVMPE-RS Multiphoton Laser Scanning Microscope equipped with a Spectra Physics dual line InSight X3 near infrared laser and 25x (1.05 NA) water-immersion objective was used to image inside intact tibial bone. Collagen autofluorescence (AF, 495-540 nm emission) and second harmonic generation (SHG, 410-455 nm emission) were collected using 880 nm excitation. Fluorescence from dTomato positive PMNs was collected using 1040 nm excitation and a 575-645 nm emission filter.

### Single cell RNA sequencing and data analysis

Murine bone marrow samples were used to generate 10X Genomics 5’ gene expression libraries which were sequenced on a NovaSeq 6000 instrument. Demultiplexing was performed with Bcl2fastq. Alignment, tagging, and gene and transcript counting were performed with Cellranger. Quality control and analysis were performed using the R package Seurat. Dead or low complexity cells were filtered by removing cells with less than 200 features and cells with a percentage of mitochondrial reads greater than 6%. Doublets were removed by filtering cells with greater than 5000 features or greater than 30,000 transcripts. Filtered data was normalized with SCTransform, and dimensional reduction was performed by UMAP. Cluster ID was performed using the R package SingleR.

### Statistics

For patient samples, per power analyses for our studies by the UNMC Biostatistics Core, a minimum of 17 patients per group was needed to detect at least a 2-fold difference in gene expression. This calculation was based assumption of a sequencing depth of 20, and a coefficient of variation of 0.4, and alpha of 0.001 to adjust for multiple comparisons for the RNA-Seq study. This sample size also achieves 80% power to detect a large Cohen’s d effect size of 1.0 at alpha=0.05 using a two-sided two-sample t-test. Additional subjects were recruited to account for any patients with low yields of PMN/working material. The continuous data were compared between groups using two-sample t test or ANOVA when appropriate. Graphpad Prism software was utilized for all other statistical analyses.

## Results

### Impact of prostate cancer progression on PMN number and viability

Blood was isolated from males from varying stages of PCa (localized/non-metastatic disease; bone metastatic hormone/castration-sensitive (CSPC); bone metastatic castration-resistant (CRPC). For comparison, blood was collected from healthy male donors. Polymorphonuclear leukocytes/neutrophils (PMNs) were isolated from blood using a negative isolation method to prevent premature activation. Although PMN numbers were similar between groups, ranging from ~1-6 million cells per mL of whole blood, there was a trend towards increased PMN numbers with progression to metastatic disease that was significantly reduced with progression from hormone sensitive metastatic disease (mCSPC) to castration resistance (mCRPC) (**Figure 1A**).

**Figure 1.**
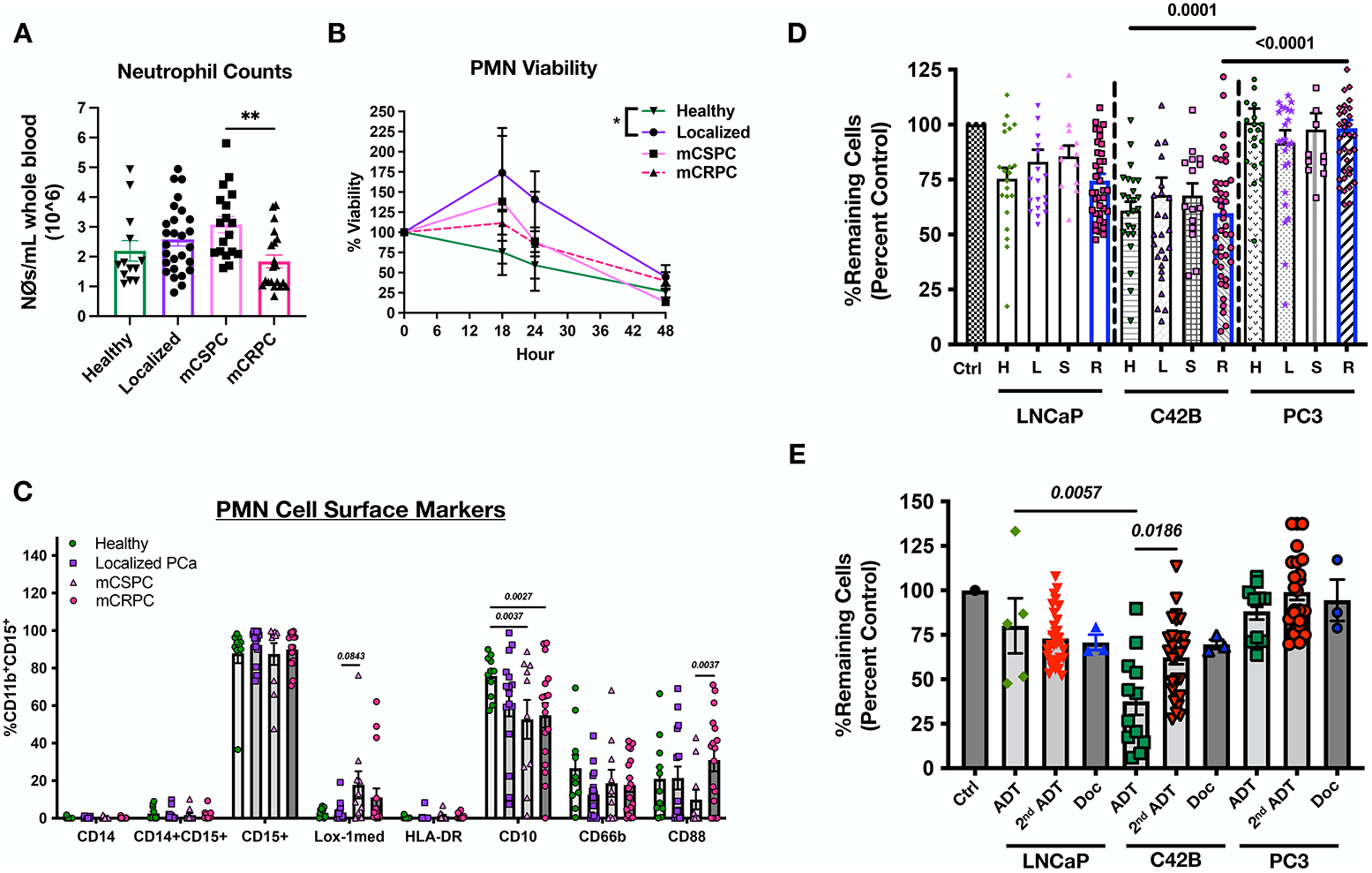
Functional and molecular analysis of peripheral blood PMNs. PMNs were isolated from blood samples using a negative isolation magnetic bead kit. (A) PMN counts per mL of blood per disease stage. One-way ANOVA was performed. **p<0.01 (B) PMN viability per specified hours post isolation using CellGro viability assay. Statistics using two-way ANOVA; statistical significance at 18 hours between healthy and localized patients; *p < 0.05 (C) FACS analysis was performed using specified cell surface markers. Plot shows percentage of marker-positive cells. Two-way ANOVA statistical analysis was performed; graph shows p-values. (D) Co-culture of patient PMNs per PCa stage with PCa cell lines (LNCaP, C42B, and PC3); 10:1 PCa to neutrophil ratio plated in triplicate. PCa cell numbers were normalized to 100% and graph shows percent remaining PCa cells after overnight culture with PMNs. H= healthy; L= localized; S= Castration-sensitive; R= Castration-resistant. (E) Co-culture of patient PMNs treatment with PCa cell lines (LNCaP, C42B, and PC3). Ctrl= Control; ADT= 1^st^ line androgen deprivation therapy; 2^nd^ Gen ADT= 2^nd^ generation androgen deprivation; Doc= docetaxel (chemotherapy).

Prior studies have demonstrated that circulating PMNs in cancer patients are predominantly immature cells and exhibit extended survival and viability, likely due to tumor-induced systemic increases in G-CSF and GM-CSF^21,22^. We examined whether PMN viability is affected by PCa disease stage. To do this, PMN viability was measured using CellTiter Glo luminescence assay, which utilizes ATP-mediated conversion of luciferin. At 18 hours post-isolation, at least 25% of PMNs from healthy men had died, 50% at 24 hours, and 75% had died by 48 hours after isolation (**Figure 1B**). This was in contrast to mouse neutrophils isolated from bone marrow, which exhibit ~70-75% cell death by 18 hours after isolation^16^. There was either extended viability or increases in cell number in the cancer groups/disease stages in the first 24 hours after isolation. Notably, PMNs from patients with localized PCa and mCSPC showed increased numbers at 18 hours (173%,155%) and 24 hours (141%, 124%), respectively. At 18 hours, post isolation, there were significantly more PMNs in blood from localized PCa in comparison to healthy men. PMNs from castration-resistant/mCRPC patients showed extended viability within the first 24 hours after isolation, with ~100% of the cells remaining at 18 hours and 75% viable at 24 hours post-isolation. By 48 hours post-isolation, ~ 25% of PMNs from healthy donors remained while there were slightly more cells in the cancer patient groups, ~30-40% viable. These results support previous studies demonstrating prolonged viability of PMNs from cancer patients compared to healthy/non-cancer patients. However, there was no difference in PMN viability between patients with mCSPC compared to patients with bone mCRPC.

### PMNs from prostate cancer patients show a combination of mature and immature myeloid cell markers

We next performed flow cytometry to gain insight into the PMN differentiation and/or activation status based upon known cell surface markers, classically found on all PMNs, and those found to distinguish immunosuppressive PMNs from pro-inflammatory, activated PMNs. PMNs were identified from CD11b^+^CD15^+^CD16^+^CD14^−^ populations. There were no differences in the percentage of CD15^+^(granulocyte marker) or CD16^+^ (marker for mature granulocytes; not shown) and there were little to no CD14^+^ (monocyte marker) cells per treatment group (**Figure 1C**). There was a slight, but not significant increase, in Lox1+ PMNs with progression to metastatic disease e.g., in comparison of localized PCa to mCSPC, indicative of an immunosuppressive phenotype. Similarly, there was a reduction in CD10^+^ PMNs in all PCa patients with tumor progression though it was most significantly reduced in patients with bone metastases (~21% reduction in HSPC and mCRPC) compared to patients with only localized PCa. CD10 expression has been associated with differentiation stage, in which its expression increases with PMN maturation, suggesting that PMN populations from PCa are less mature and immunosuppressive. Even still, ~50% of PMNs from all stages expressed CD10 and fewer than ~20% were Lox-1 positive. There was a slight reduction in CD66b, a marker of activated PMNs. Interestingly, there was a reduction in PMNs expressing CD88, the compliment C5a receptor, in progression from localized PCa to mCSPC; however, it was significantly increased on PMNs from mCRPC suggesting a transition in activation with the change in therapeutic response. These data highlight disease-specific changes in PMN cell surface markers that may dictate PMN function throughout PCa progression.

### PCa progression induces a pro-inflammatory PMN gene signature

To gain further insight into molecular changes of PMNs, we performed bulk RNA sequencing on PMNs per disease stage (n=16 total (4/healthy; 4/localized; 3/mCSPC; 5/mCRPC); note: we initially intended to analyze 4 per group however, response to androgen therapy changed in one CSPC patient e.g., the patient became resistant to androgen therapy after grouping by diagnosis but prior to PMN isolation and so there were 3 mCRPC and 5 mCRPC samples sequenced. Because PMNs are terminally differentiated cells, there was a low total number of genes altered per group. However, there were distinct gene changes per stage of disease. Based on the QC plot, there were >10,000 genes detected per treatment group out of approximately 25,000 genes analyzed (full gene list can be found at GEO Accession: GSE197609) (**Figure 2A**). After controlling for the false discovery rate, we found that overall, there were more genes turned “on” than “off” throughout PCa progression. When comparing disease stage, the most PMN genes were altered when comparing mCRPC patients to healthy individuals; genes up-/down-regulated by more than 2-fold, (*p*<0.05: 135 up/22 down; *p*<0.01: 44 up/11 down) demonstrating that there is a significant change in the molecular profile of PMNs in PCa patients (**Figure 2A**). However, due to the variations in clinical regimen of the sequenced patients, there were no identified treatment-associated genetic signatures (**Figure 2B**). When comparing PMNs from healthy men compared to CRPC patients, cancer-derived PMNs exhibit a pro-inflammatory molecular profile, including increased expression of IL-8, G-CSF receptor, Lactoferrin (LFN), MMP8, and CD177 (**Figure 2C**). There was also a significant increase in arginase I (ARG1), which has been typically associated with immunosuppression^23^. Significantly altered genes in mCRPC, compared to healthy men, were further analyzed by Ingenuity Pathway Analysis (IPA) software (**Supp Figure 1A**). The top 2 canonical pathways identified from gene enrichment analysis were: 1) glycolysis I (5 of 26 genes), and 2) 6-Phosphofructo-2-Kinase/Fructose-2,6-Bisphosphatase 4 (PFKFB4) signaling pathway (6 of 46 genes) both which are associated with altered redox signaling and glucose metabolism^24^; these findings are in support of our previous findings where we identified that BM-PCa alters redox metabolism genes in bone marrow neutrophils and increases neutrophils reactive oxygen species production (ROS)^25^.

**Figure 2.**
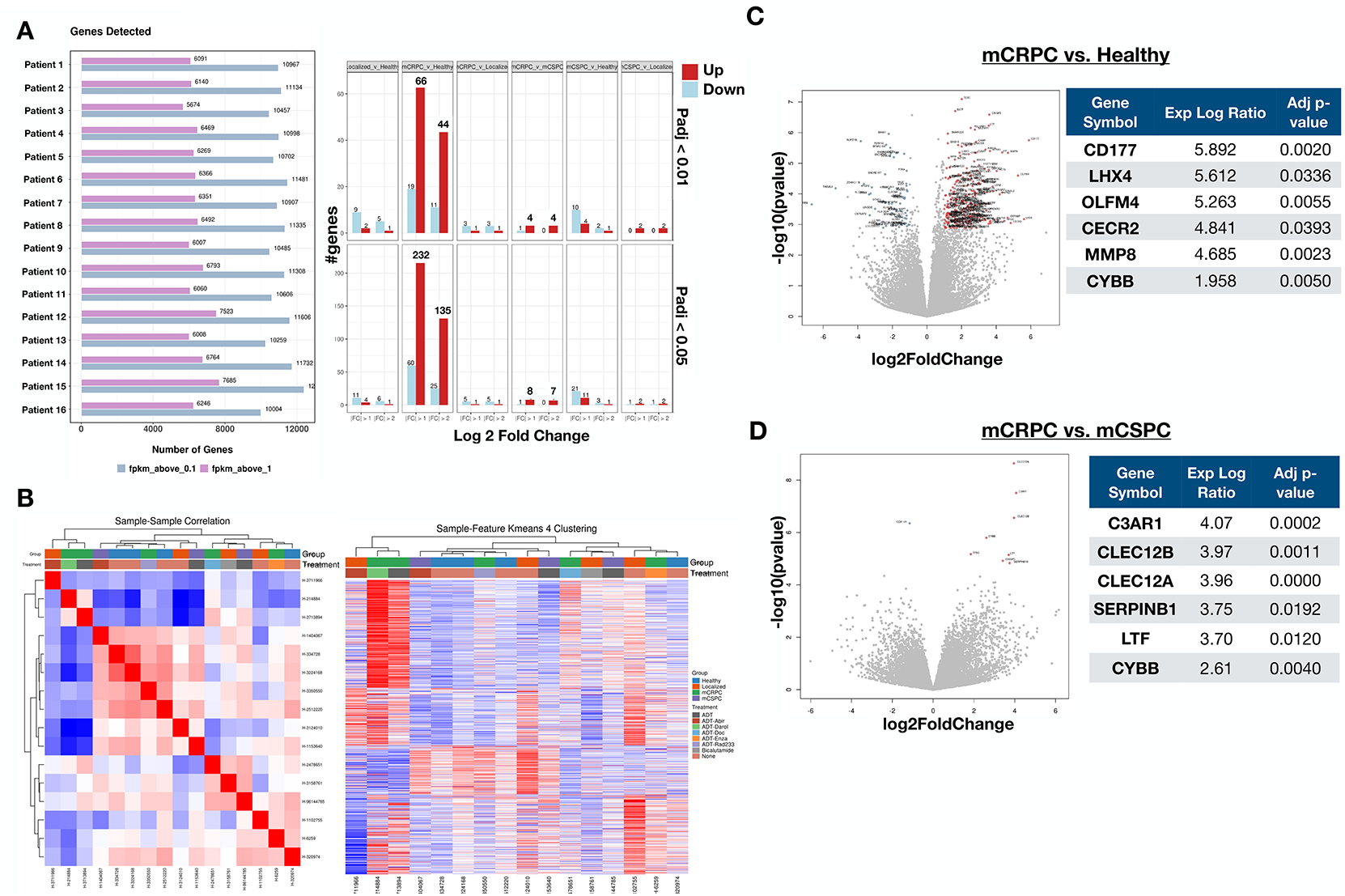
RNA sequencing analysis of PCa-derived PMNs. Bulk RNA sequencing of blood PMNs was performed on an Illumina NovaSeq with paired-end 100bp reads (n per group=4/healthy; 4/localized; 5/mCSPC; 3/mCRPC). (A) Left, plot shows number of genes dictated per patient per fkpm. Right, number of genes significantly up-regulated (red) or down-regulated (blue) per disease stage group. (B) Left, heat map of sample-to-sample correlation of all patients; right, heatmap of K means clustering of all patients with PCa stage and treatment. (C) Volcano plot of mCRPC compared to healthy males altered genes. Table shows most up-regulated genes in mCRPC. (D) Volcano plot of differentially altered genes from mCRPC compared to mCSPC patients. Table shows most up-regulated genes in mCRPC.

For all other group comparisons, there were fewer than 10 genes up- or down-regulated more than 2-fold. The fewest genetic changes occurred in PMNs during the transition from localized PCa to metastatic disease (localized vs. mCSPC). There were only 3 genes in total altered in this comparison (up/down by more than 2-fold (p<0.01: 2/0; p<0.05: 2/1)), suggesting that the least molecular change in PMNs of patients that progressed from localized PCa to the development of metastases (**Figure 2A; Supp Figure 1**). Surprisingly, there were more PMN genes significantly up-regulated when comparing hormone-responsive mPCa compared to mCRPC (genes up more than 2-fold, (p<0.01: 4; p<0.05: 7; there were no genes down-regulated)), suggesting that PCa acquired resistance to hormone-targeted therapy significantly impacts the PMN transcriptome. There were some shared upregulated genes, but no shared downregulated genes, when examining our comparisons of mCRPC vs. healthy and mCRPC vs. mCSPC, including CYBB (**Figure 2D; Supp Figure 1B & 1C**). However, there were some genes that were only altered with progression to castration-resistant disease, including complement 3 receptor 1 (C3AR1) and C-type Lectin Receptors 12A & 12B (CLEC12A/12B), which are all major inducers of inflammation suggesting a stage-specific shift to a highly pro-inflammatory PMN population (**Figure 2D**).

We previously observed an increase in CYBB expression in human bone marrow PMNs upon treatment with BM-PCa conditioned media (C42B), compared isogenic non-metastatic PCa (LNCaP)^25^. We further identified an increased release in reactive oxygen species and activation of antioxidant signaling pathways from PMNs after direct culture with PCa^25^. Thus, we examined antioxidant protein expression in patient-derived PMNs (n=8/group). We noticed there was variability in the protein content but also changes in many of the housekeeping genes, which also notably are metabolic enzymes e.g., GAPDH (**Supp Figure 2A**, Ponceau stain), preventing densitometry normalization. Thus, we quantified overall positivity of specific antioxidants, superoxide dismutases (SODs) and catalase. Similar to our previous findings, we did see altered expression of SOD1 with PCa progression to metastasis and therapy-resistant disease (**Supp Figure 2B**) but no little to no change in other antioxidants. Specifically, there were fewer SOD1-expressing PMNs in patients with metastatic diseases (~2-3 of 8 patients compared to 6 of 8 patients from localized PCa and healthy men). This suggests there may be a shift in the redox metabolism of PMNs from metastatic patients.

### Second generation hormone therapy abrogates neutrophil cytotoxicity

In preclinical PCa studies, our lab identified that bone marrow neutrophils are protective against PCa growth in bone. However, neutrophil cytotoxicity was suppressed as the tumor became resistant and progressed within the bone compartment. Based on these findings, we investigated the cytotoxic potential of peripheral blood PMNs in PCa patients. To do this, isolated patient PMNs (n= >17/group) were cultured overnight in a 10:1 PMN:PCa ratio with PCa cells (non-metastatic LNCaP, C42B (bone metastatic LNCaP cells), and PC3 (bone metastatic, AR-negative, PMN-resistant)). For these analyses, we excluded PMNs isolated from PCa patients with observable visceral metastases. After overnight culture, the remaining cancer cells were counted via Trypan Blue exclusion assay, as described^16^. Patient PMNs from all disease stages significantly induced average cell death of LNCaP (~25%) and C42B (~35-40%). Surprisingly, PMNs from CRPC patients (bar graph outline in blue, **Figure 1D**) demonstrated significant killing of both LNCaP and C42B cell lines. However, there was a range of cell death of LNCaP (~25-50%) and even more robust killing of C42B with a wider range (~0-95% induced cell death). In contrast, PC3 cells, which are AR-negative, were resistant to PMNs from all disease stages similar to what we had previously shown^16^ and in some cases PMNs promoted proliferation of PC3 cells (**Figure 1D**). PMNs from healthy patients were cytotoxic to LNCaP and C42B, killing 25% and ~30%, respectively. However, PC3 were resistant to PMNs from even healthy individuals. *These findings support the fact that there are cancer intrinsic factors that affect neutrophil killing*. Patient PMNs were most cytotoxic towards the AR-positive PCa cells, however there was some variability in killing capacity that appeared to be dependent on treatment.

We next categorized the cytotoxicity data by therapy. The majority of all PCa patients in this study received initial ADT, i.e., Leuprolide, an antagonist of gonadotrophin-releasing hormone, while patients whose disease progressed (as determined by rising PSA levels) were administered 2^nd^ generation non-steroidal anti-androgens (including abiraterone or enzalutamide). Based on the diversity of ARSI treatments in our patient cohorts, we compared PMN cytotoxicity from PCa patients who had only received initial ADT (those who had only received Luprolide or orchiectomy) at the time of blood collection with those patients who received 2^nd^ generation/additional ADT, including ARSIs. All PCa stages were represented in both cohorts of 1^st^ line ADT vs. 2^nd^ gen ADT, including those with localized and metastatic PCa. We also included one patient who received chemotherapy (Docetaxal (Doc)). Interestingly, we found that PMNs from patients treated with androgen deprivation therapy (ADT) alone, were the most cytotoxic to C42B, killing ~75% of C42B. However, PMNs from patients who received ADT combined with a 2^nd^ generation androgen therapy (such as Enzalutamide or Abiraterone) showed impaired cytotoxicity (**Figure 1E**). For an additional BM-PCa cell line for comparison, we also examined PMN killing of 22Rv1 (which were derived from CWR22 xenograft^26^, which are bone metastatic and express the constitutively active AR variant V7. Of 3 patients tested, 1 patient which received ADT + 2^nd^ Gen ADT (noted by the top 3 co-culture points/replicates), showed an inability to kill 22Rv1 compared to the other 2 patients that received only ADT (**Supp Figure 3**). In total, PMNs from patients treated with ADT in combination with 2^nd^ generation androgen inhibition were significantly less cytotoxic than ADT alone. These data suggest that androgen regulation is a critical mediator of anti-tumor PMN response. Next, we examined the therapeutic value of this hypothesis in an *in vivo* intratibial metastasis model.

**Figure 3.**
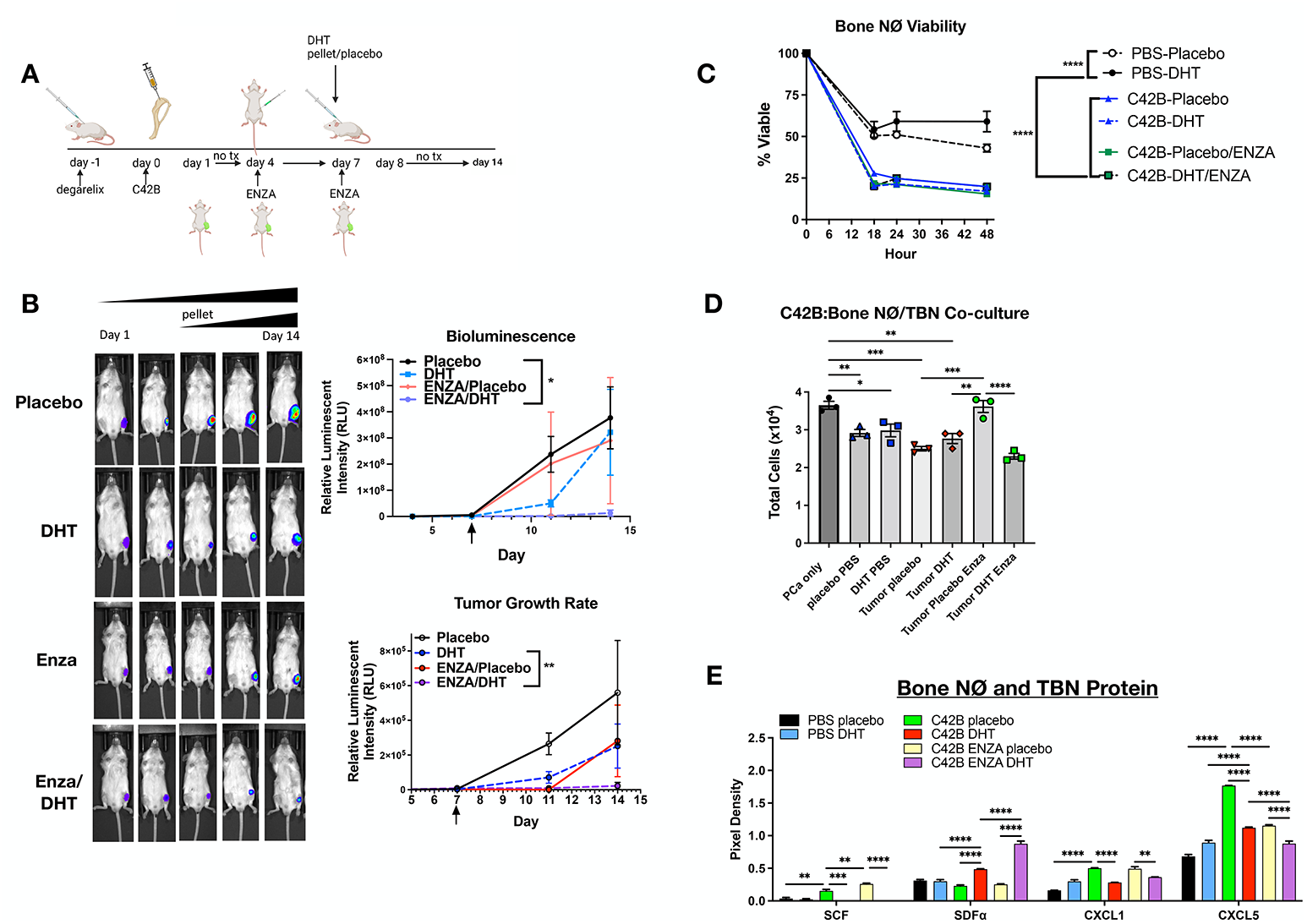
Bipolar androgen therapy (BAT) and ARSI regulation of PMN cytotoxicity. (A) Schematic of pre-clinical trial; generated by Biorender.com. Standard-care prostate cancer treatment regimen was applied in a preclinical trial of BAT and ARSI in male SCID Beige mice. Mice were chemically castrated using Degarelix and tibia were inoculated with luciferase-expressing C42B cells. One week post tumor inoculation, mice were treated with enzalutamide (Enza) (treated weekly for the entire study) and the following week were subcutaneously implanted with either slow-release dihydrotestosterone (DHT) or placebo pellets. (B) Quantitative bioluminescence imaging. Top graph shows quantitation of relative luminescent intensity (RLU) in photons/cm^2^/second over time. Bottom graph shows RLU normalized to bioluminescence at Day 1 post tumor inoculation. Arrows point to beginning of ADT treatment. (C) Real Time Glo MT cell viability assay of tumor-bone neutrophils (TBNs) isolated from tumor-bone at the end of study, day 14. Two-way ANOVA statistical analysis performed; ****p<0.0001 at 18 hours post-isolation. (D) C42B ex vivo co-culture with TBNs from each mouse group: placebo PBS (saline-injected mice with placebo pellet), DHT PBS (saline-injected mice with DHT pellet), tumor placebo (C42B-injected mice with placebo pellet), tumor DHT (C42B-injected mice with DHT pellet), tumor placebo Enza (C42B-injected, enzalutamide-treated, placebo pellet), and tumor DHT Enza ((C42B-injected, enzalutamide-treated, DHT pellet). For co-culture, C42B were plated in triplicate, cultured with TBNs overnight, and remaining C42B counted using Trypan Blue exclusion assay. Graph shows C42B numbers after culture with TBNs. (E) Cytokine array of TBNs. Two-ANOVA statistical analysis was performed. **p<0.01, ***p<0.001, ***p<0.0001.

### Bipolar androgen therapy manages BM-PCa progression and restores neutrophil immune response

Disease recurrence in PCa is typically associated with acquired resistance to androgen therapy. mPCa is treated with ARSIs upon disease recurrence; however, the tumor inevitably adapts to reduced AR signaling. A previous clinical study leveraged the dynamic tumor through applied high dose testosterone to “shock” the tumor by overloading it with more testosterone than feasible to utilize, followed by cycles of ARSIs and testosterone to stabilize tumor progression, known as bipolar androgen therapy (BAT)^27,28^. Based on our findings demonstrating ARSI regulation of patient PMN cytotoxicity, we tested the potential for androgen regulation using BAT as an approach to enhance neutrophil immune response *in vivo*. To do this, male mice were first chemically castrated using Degarelix^29,30^. Mice then received an intratibial injection of luciferase-expressing C42B cells (n=4-5/group). To mimic a standard-of-care clinical regimen, mice were treated with either enzalutamide or a vehicle control and, one week later, subcutaneously implanted with a 21-day slow-release dihydrotestosterone (DHT) pellet, or placebo (**Figure 3A**). Surprisingly, we found that DHT alone suppressed tumor burden, demonstrated by reduced bioluminescent intensity (**Figure 3B**). Enzalutamide suppressed tumor burden more than DHT alone; however, combined DHT and Enzalutamide had the most significant impact on tumor burden compared to the control/placebo group. Although enzalutamide suppressed tumor burden, the tumor growth rate appeared to be nearly identical when comparing sole treatment of DHT or Enzalutamide. Combination DHT and Enzalutamide significantly suppressed tumor growth rate compared to DHT alone (**Figure 3B**).

We next isolated neutrophils from the tumor bones for analyses of function, in comparison to neutrophils from saline-injected bones. Neutrophils from non-tumor limbs treated with DHT were the most viable, 75% were still viable in serum-free media 48 hours after isolation compared to 50% from placebo-treated mice and only ~25% remaining viable cells from all tumor mice (**Figure 3C**). Tumor-bone neutrophils (TBNs) and tumor-naïve neutrophils (from saline-injected tibia) were co-cultured ex vivo with C42B to examine the impact of treatment on TBN cytotoxicity. Neutrophils from both PBS and DHT-treated non-tumor bearing mice induced ~50% cell death. Similar to PMNs from patients treated with 2^nd^ Gen ADT, cytotoxicity of TBNs from enzalutamide-treated (Enza-placebo group) mice was completely diminished. However, the addition of DHT restored TBN cytotoxicity when combined with Enzalutamide (**Figure 3D**). These data suggest that androgen regulation can be critically leveraged to enhance neutrophil anti-tumor immune response.

Overall, there were very few changes in the percentages of isolated bone marrow myeloid cells though there were notably reduced myeloid cell numbers ((CD11b^+^, CD45^+^) and (CD11b^+^, Ly6C^+^/Ly6G^+^)) in enzalutamide-treated mice, revealed by FACS (**Supp Figure 4A**). We previously showed that the Ly6C/Ly6G bone marrow population are predominantly neutrophil precursor cells that exhibit cytotoxicity against prostate cancer and suppression of T cell proliferation^16^. Upon examining TBN-derived proteins via protein array, we found that there were few changes in myeloid-regulating growth factors, such as G-CSF and GM-CSF. However, there was significantly more M-CSF expressed in TBNs from Enza/DHT mice compared to DHT-treated mice (**Supp Figure 4B**). TBNs from mice treated with Enza and DHT showed increased TNFalpha and TNF receptor expression which supported the cytotoxic and reactive response against C42B identified in *ex vivo* analyses (**Supp Figure 4B**). Interestingly, in comparison to placebo-treated mice, there was little to no expression of Stem Cell Factor (SCF), the neutrophil differentiation marker, in neutrophils and TBNs from all DHT-treated groups including tumor naïve, tumor-bearing DHT-treated and tumor-bearing mice treated with combinational DHT and enzalutamide (**Figure 3E**). SCF levels were significantly lower than the other treatment groups suggesting that SCF regulation may be important in androgen regulation of TAN function. These findings suggest that androgens play a critical role in neutrophil ability to kill PCa.

**Figure 4.**
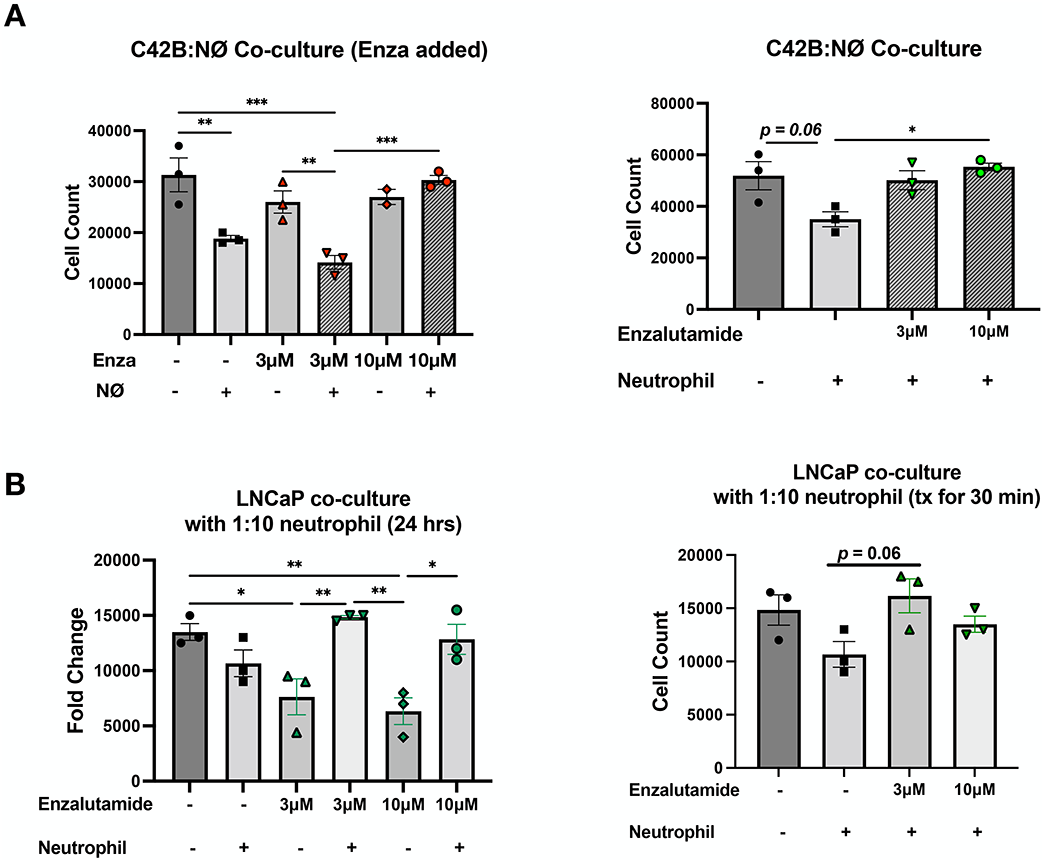
Enzalutamide directly suppresses neutrophil function and cytokine ex vivo. (A) Left, C42B direct co-culture with primary mouse bone marrow neutrophils. Enzalutamide (Enza) was added to the culture when neutrophils were seeded. Right, primary mouse bone marrow neutrophils were treated with enzalutamide for 30 minutes and then added to C42B for direct co-culture overnight. Graphs show C42B cells after overnight culture via Trypan Blue exclusion assay. (B) Left, LNCaP direct co-culture with mouse neutrophils and enzalutamide added directly to culture. Right, neutrophils were treated with enzalutamide for 30 minutes before addition to culture. Graphs show C42B cells after overnight culture via Trypan Blue exclusion assay. One way ANOVA statistical analysis was performed; *p<0.05, **p<0.01, ***p<0.001.

### Androgen signaling regulates neutrophil cytotoxicity, viability, migration and NETosis

To further test this hypothesis, C42B cells were co-cultured with bone marrow neutrophils isolated from C57BL/6 mice, in the presence of enzalutamide at 2 different doses (high dose-10 μM, low dose-3 μM) for 24 hours. As previously seen, mouse neutrophils significantly killed approximately 45% of the C42B cells i.e., there were 45% fewer C42B after culture with neutrophils and we previously determined that this was due to neutrophil induction of apoptosis of prostate cancer cells. However, high dose enzalutamide completely abrogated neutrophil killing of cancer cells (**Figure 4A, left**). To delineate which cellular component was most affected by enzalutamide, neutrophils were pre-treated with enzalutamide for 30 minutes before culturing them with C42B for 24 hours. Surprisingly, neutrophils pre-treated with low and high dose enzalutamide prior to culture with C42B were unable to induce cell death of C42B (**Figure 4A, right**). These findings suggest that enzalutamide directly inhibits neutrophil cytotoxicity towards prostate cancer. This phenomenon was also seen in neutrophil-LNCaP cell cultures (**Figure 4B**) demonstrating that ARSI in neutrophils reverses their cytotoxicity against both nonmetastatic and metastatic PCa.

Previous studies have linked neutrophil function to specific tumor-associated phenotypes^31^. This includes a prolonged viability of tumor-associated neutrophils, a phenotype typically associated with immature/undifferentiated populations. To test the role of enzalutamide on neutrophil viability, primary mouse bone marrow neutrophils were treated for 2 hours with C42B-conditioned media supplemented with enzalutamide (3 μM and 10 μM); viability was measured for 24 hours. Compared to control (DMSO)-treated neutrophils, both doses of enzalutamide prolonged neutrophil viability over time. There was an increase in luminescence of low dose-enzalutamide treated neutrophils in the first 6 hours suggesting an increase in neutrophil numbers i.e., proliferation. High dose enzalutamide prolonged neutrophil viability and increased proliferation (**Supp Figure 5A, 5B**); these results were confirmed using Trypan Blue exclusion assay of enzalutamide-treated neutrophils. These findings demonstrate that enzalutamide increases neutrophil proliferation and viability, though it also inhibits neutrophil killing of PCa cells.

**Figure 5.**
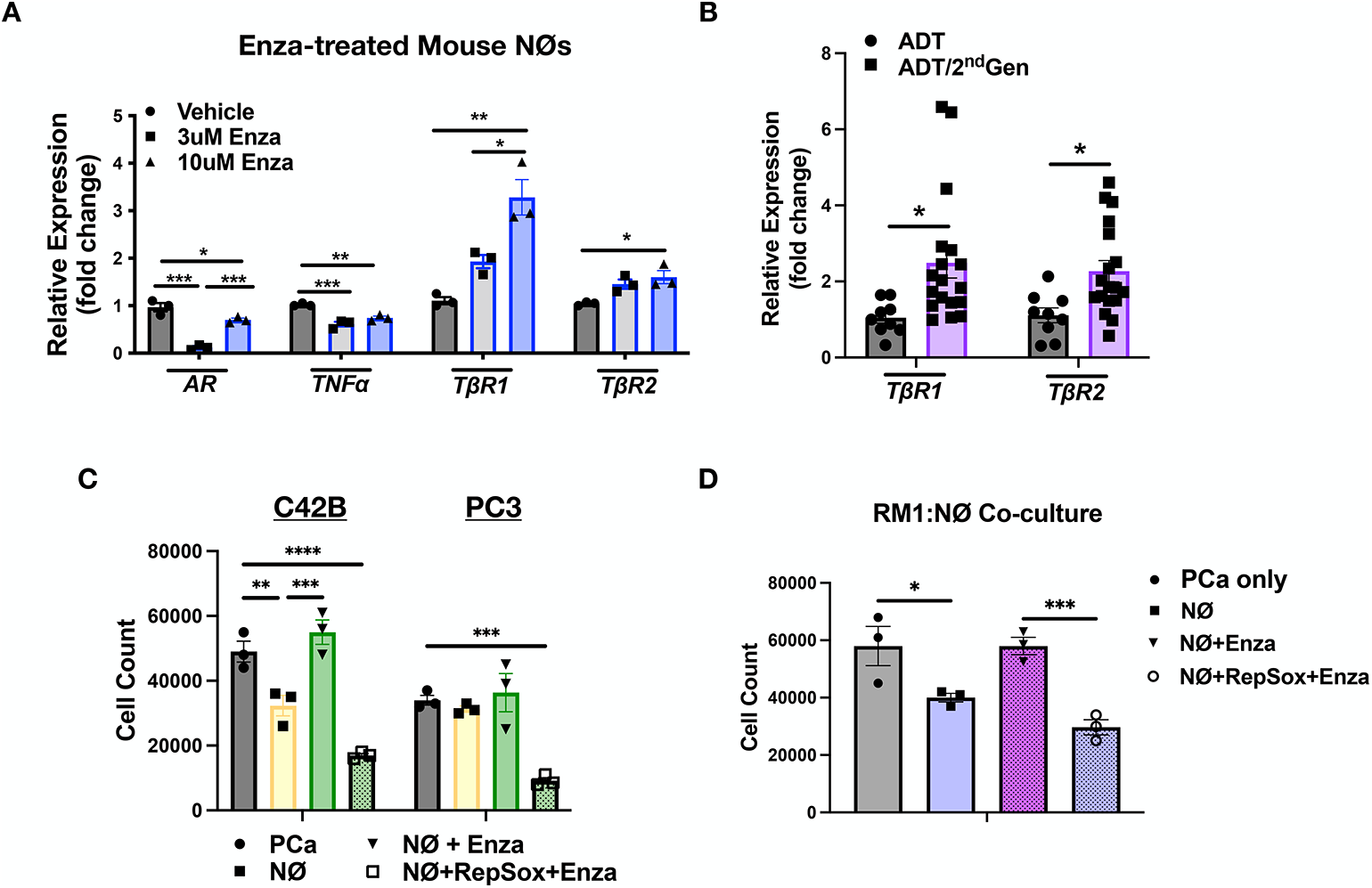
ARSI suppresses neutrophil anti-tumor response via regulation of TβR1 expression. (A) RealTime quantitative PCR (RT-qPCR) of mouse primary bone marrow neutrophils treated with enzalutamide. Genes profiled: androgen receptor (AR), tumor necrosis factor alpha (TNFa), transforming growth factor beta receptor 1 (TβR1). (B) RT-qPCR of human patient PMNs treated with either 1st line androgen deprivation therapy (ADT) or 2nd line ADT (includes all ARSIs plus 1st line ADT). (C) C42B and PC3 co-culture assay with mouse bone marrow neutrophils pre-treated for 30 minutes with either RepSox (small molecule kinase inhibitor of TβR1; 5nM), Enzalutamide (3 μM) or a combination prior to addition to overnight culture. Graph shows PCa cell counts after overnight culture. (D) Mouse RM1 PCa culture with mouse neutrophils pre-treated with RepSox and/or Enzalutamide for 30 minutes prior to addition to culture. Graph shows RM1 cell counts after overnight culture. One-way or two-way ANOVA statistical analysis were performed where appropriate. *p<0.05, **p<0.01, ***p<0.001, ****p<0.0001.

Based on our previous findings of PCa-mediated neutrophil function, we next examined the impact of androgen inhibition on neutrophil migration. We previously showed that neutrophils migrate to PCa-conditioned media, independently of CXCR2 signaling^16^. To determine whether androgen signaling is important for neutrophil migration, mouse bone marrow neutrophils were treated with enzalutamide or, for comparison, abiraterone, which inhibits CYP17A1^32^, an enzyme important for androgen synthesis. Migration was examined using modified Boyden Chamber assay. As previously shown by our group, neutrophils migrated to C42B soluble factors significantly more (~5-fold) than to RPMI alone. Interestingly, enzalutamide further enhanced neutrophil migration to C42B media (7-fold compared to control) whereas abiraterone had little to no impact on neutrophil migration to C42B factors (**Supp Figure 5C**). This suggests that androgen signaling through neutrophil AR regulates migration toward BM-PCa.

Our lab previously demonstrated that PCa induces release of NETs, such that the highest number of NETs was produced by neutrophils treated with BM-PCa media compared to non-metastatic PCa and nonmaligant prostate epithelial cells. To examine whether NETosis is regulated by androgen signaling, bone marrow neutrophils were treated with C42B-conditioned media supplemented with either enzalutamide or abiraterone for comparison to conditioned media alone or the positive control, base RPMI media supplemented with PMA to induce NET formation. As previously seen, C42B enhanced NET production (2-fold); however, enzalutamide stimulated increased NETosis (4-fold) (**Supp Figure 5C**). Similar to the migration assay, abiraterone had no impact on C42B-induced NETosis. These findings collectively suggest that neutrophil AR is an important mediator of PCa-induced neutrophil migration and NETosis.

### Second generation AR therapy regulates neutrophil gene expression *in vitro* and in patients

Based on current standard treatment for BM-PCa patients, we treated mouse bone marrow neutrophils with enzalutamide and measured viability and proliferation. Enzalutamide treatment increased neutrophil viability and cell numbers, characteristics of immature neutrophils. We next examined enzalutamide-mediated gene regulation potentially contributing to neutrophil function. Primary mouse bone marrow neutrophils were treated with 2 doses of enzalutamide (3 μM and 10 μM) for 2 hours and RNA isolated using Trizol for downstream RealTime qPCR analysis with a focus on genes previously identified to regulate neutrophil response, TNFalpha, and TGFβ receptors^15^. We additionally measured neutrophil AR expression and interestingly found that enzalutamide significantly reduced neutrophil AR expression (3uM-~9-fold; 10um-~2-fold) compared to DMSO-treatment (**Figure 5A**). There was also a significant reduction in expression of inflammatory gene, TNFalpha gene (2-fold reduction). To our surprise, there was a dose-dependent increase in the serine-threonine kinase, Type 1 TGFβ Receptor (R) of enzalutamide-treated neutrophils (**Figure 5A**) and a slight, but significant, increase in the Type 2 TGFβ receptor, which binds TGFβ ligands.

For comparison, we examined expression of TGFβ receptors in patient blood-derived neutrophils from patients who received AR therapy. PMNs from patients treated with ADT and 2^nd^ generation ADT, showed significantly increased TβR2 and TβRI gene expression (~2-fold) compared to patients treated with ADT alone (**Figure 5B**). These findings suggest that AR regulates TGFβ signaling in neutrophils which may be significant in the regulation of neutrophil function.

### Targeted TβRI inhibition overcomes enzalutamide-mediated suppression of function

Based on the identified anti-inflammatory and immunosuppressive roles of TGFβ, we next tested the importance of neutrophil TβRI expression on neutrophil function. Specifically, primary bone marrow neutrophils were treated with RepSox, a small molecule kinase inhibitor of TΒRI, or vehicle control, and then subsequently cultured them with C42B and PC3 for comparison. Notably, inhibition of neutrophil TβRI, reversed the enzalutamide suppression of neutrophil cytotoxicity while significantly enhancing neutrophil-induced C42B cell death even more than the absence of enzalutamide, inducing 75% C42B cell death compared to 50% with no enzalutamide (**Figure 5C**). Similarly, TβRI-inhibited neutrophils induced cell death of PC3 which are notably resistant to neutrophil-induced cell killing. This phenomenon was also verified using mouse RM1 prostate cancer cells where RepSox reversed Enzalutamide suppression of neutrophils and demonstrating that these findings are conserved between species (**Figure 5D**).

TGFβ is characteristically a pleiotropic cytokine utilized by multiple cells within the bone marrow, which complicates therapeutic intervention. To gain insight into TβRI expression in mouse bone marrow and the potential for TβRI-focused therapy, single cell RNA sequencing data from male C57BL/6 bone marrow was analyzed with a specific focusing on TβRI expression in Ly6G+ cells and overall TβRI expression in bone marrow cells (**Supp Figure 6A**). There was TβRI expression in bone marrow neutrophils, and specifically, all neutrophil subpopulations within the bone marrow (**Supp Figure 6A**). However, TβRI was not expressed in all Ly6G+ cells i.e., granulocytes, suggesting that TβRI expression may be more relevant in the instance of androgen regulation. To gain insight into TβRI expression in PCa metastases, we performed fluorescent immunohistochemistry on matched liver and bone metastasis from prostate cancer patients (n=3; all who received 2^nd^ Gen ADT), to identify TβRI^+^ neutrophils, identified by myeloperoxidase (MPO). In bone metastasis samples, we found co-localization of TβRI and MPO (in 3 of 3 patients) whereas there was little to no co-localization of TβRI and MPO in the matched liver metastases (co-localization was seen in 1 of 3 patients) suggesting that TβRI-expressing neutrophils may be more conserved in bone metastases, compared to liver metastasis. Further, analysis of patient PMN genes from our study revealed TGFβ to be a major upstream mediator of gene regulation in mCRPC patient PMNs compared to healthy men and a critical regulator of TβRI expression **(Supp Figure 6B)**.

**Figure 6.**
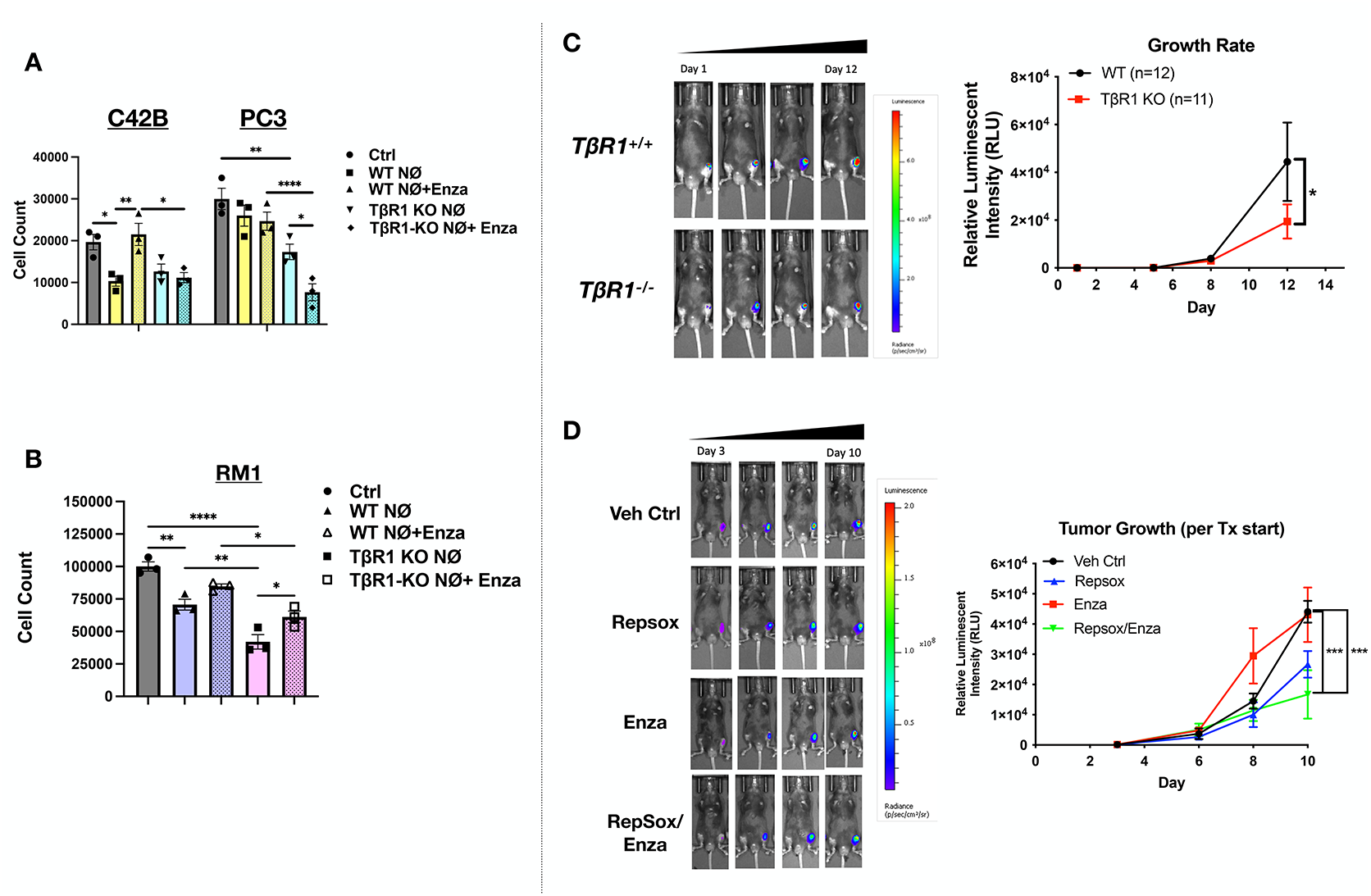
Genetic deletion of TβRI and combined ARSI and TβRI therapy suppresses mCRPC progression. (A) C42B and PC3 co-culture assay with wildtype TβRI and TβRI-null mouse bone marrow neutrophils pre-treated with Enzalutamide (3 μM), where shown, prior to addition to overnight culture. Graph shows PCa cell counts after overnight culture. Two-way ANOVA statistical analysis was performed. *p<0.05, **p<0.01, ****p<0.0001. (B) RM1 co-culture assay with wildtype TβRI and TβRI-null mouse bone marrow neutrophils pre-treated with Enzalutamide (3 μM), where shown, prior to addition to overnight culture. Graph shows PCa cell counts after overnight culture. One-way ANOVA statistical analysis was performed. (C) RM1 intratibial bone metastasis model in TβRI KO mice. Luciferase-expressing RM1 cells were injected into tibia of 6-7 week male wildtype TβRI (n=12) or neutrophil-selective TβRI-null mice (n=11) and tumor burden measured longitudinally for the entire study using bioluminescence imaging. Representative image (left) and bioluminescence quantitation (right, photons/cm2/sec). Two-way ANOVA statistical analysis was performed. *p<0.05 at Day 12 of the study. (D) RM1 intratibial metastasis in preclinical therapeutic trial. Luciferase-expressing RM1 was injected intratibially in 6-week C57BL/6 mice (n=5-7) and randomized via bioluminescence into 4 treatment groups: 1) vehicle control (DMSO), 2) RepSox (5mg/kg), 3) Enzalutamide (Enza; 10mg/kg), 4) RepSox combined with Enza. Representative images (left) and quantitation (right). Two-way ANOVA statistical analysis was performed; graph shows significance at end of study, at Day 10.

Based on this evidence, we generated a neutrophil-selective TβRI knockout mouse model using the Catchup mouse model, which express Cre-recombinase and dTomato under the Ly6G locus (**Supp Figure 8A, left**). Catchup mice were crossed with TβRI floxed mice and backcrossed onto C57BL/6 mice for a minimum of 6 generations. Protein analysis of multiple mouse tissues revealed a neutrophil-specific loss of TβRI with no change in other tissues (**Supp Figure 8A, right**). There were no differences in neutrophil cell number with genetic deletion of TβRI (**Supp Fig 8B**), however there were significantly fewer myeloid cells (CD45^+^CD11b^+^) in the bone marrow (**Supp Figure 8C**). Although there appeared to be fewer Ly6G^+^ cells, this was not surprising considering that the Ly6G knock-in in the Catchup mice disrupts Ly6G expression (**Supp Figure 8C**) without impacting cellular function^17^. Surprisingly, there was a significant increase in CD19^+^ B cells, and more CD8^++^ cells in TΒRI^−/−^ (**Supp Figure 8C**) suggesting an indirect impact of neutrophil TGFB signaling on other cell populations.

Next, we examined killing capacity of TβRI knockout neutrophils against BM-PCa cells (C42B, PC3, and 22Rv1). Neutrophils induced death of C42B, independently of TβRI expression; however, TβRI knockout neutrophils induced cell death of PC3, neutrophil-resistant (**Supp Figure 8D**). Neutrophils also induced 22Rv1 death, but there was little additional impact in the absence of TβRI (**Supp Figure 8D**). To examine the impact of neutrophil-selective TβRI in the context of androgen-mediated activity, we isolated primary mouse bone marrow neutrophils from TβRI-null and TΒRI floxed mice, treated them with low-dose enzalutamide (3 μM) or vehicle for 30 minutes and cultured them overnight with C42B and PC3 cells. As seen in our previous experiments, enzalutamide completely suppressed wildtype neutrophil killing of C42B; however, TβRI-null neutrophils sufficiently killed ~50% of C42B cells even after enzalutamide treatment. Even untreated TβRI-null neutrophils killed ~40% of PC3 cells, but killed ~75% of PC3 after treatment with Enzalutamide further suggesting that TβRI inhibition may activate neutrophils against androgen-insensitive metastatic prostate cancer (**Figure 6A**) and is even further enhanced in the presence of ARSIs. For comparison, mouse RM1 prostate cancer cells were cultured with TΒRI-null neutrophils in the presence of enzalutamide. Similar to human PCa, enzalutamide suppressed neutrophil-induced RM1 death and this phenomenon was significantly reversed by genetic TΒRI inhibition in neutrophils (**Figure 6B**).

Last, we sought to determine whether these data translated to the bone microenvironment and utilized intratibial bone metastasis models to test the impact of neutrophil selective TΒRI loss *in vivo*. Luciferase-expressing RM1 cells were injected into tibia of TβRI^fl/fl^ and TβRI^−/−^ mice. Based on bioluminescent changes, loss of TβRI in neutrophils alone significantly suppressed RM1 growth in bone (**Figure 6C**). Despite these findings, it would be difficult to target TβRI inhibition solely to neutrophils since it is expressed by other cell populations in the bone microenvironment. Thus, we tested a more clinically feasible therapeutic approach and examined the impact of systemic inhibition of TβRI and AR single or in combination in BM-PCa. We found that RM1 in vivo did not respond to enzalutamide and represented a more aggressive cancer. TβRI inhibition alone appeared to suppress prostate tumor growth in bone but there was an additive effect on tumor burden in the presence of enzalutamide (**Figure 6D**).

Collectively, these findings strongly support the conclusion that androgen deprivation, specifically AR inhibition, increases neutrophil TβRI and suppresses their cytotoxicity against prostate cancer (**Figure 7**). Pharmacologic and genetic inhibition of neutrophil TβRI completely rescued this phenomenon. Thus, TβRI inhibition can be used to overcome neutrophil suppression and to simultaneously target both AR-responsive and AR-insensitive bone metastatic prostate cancer.

**Figure 7.**
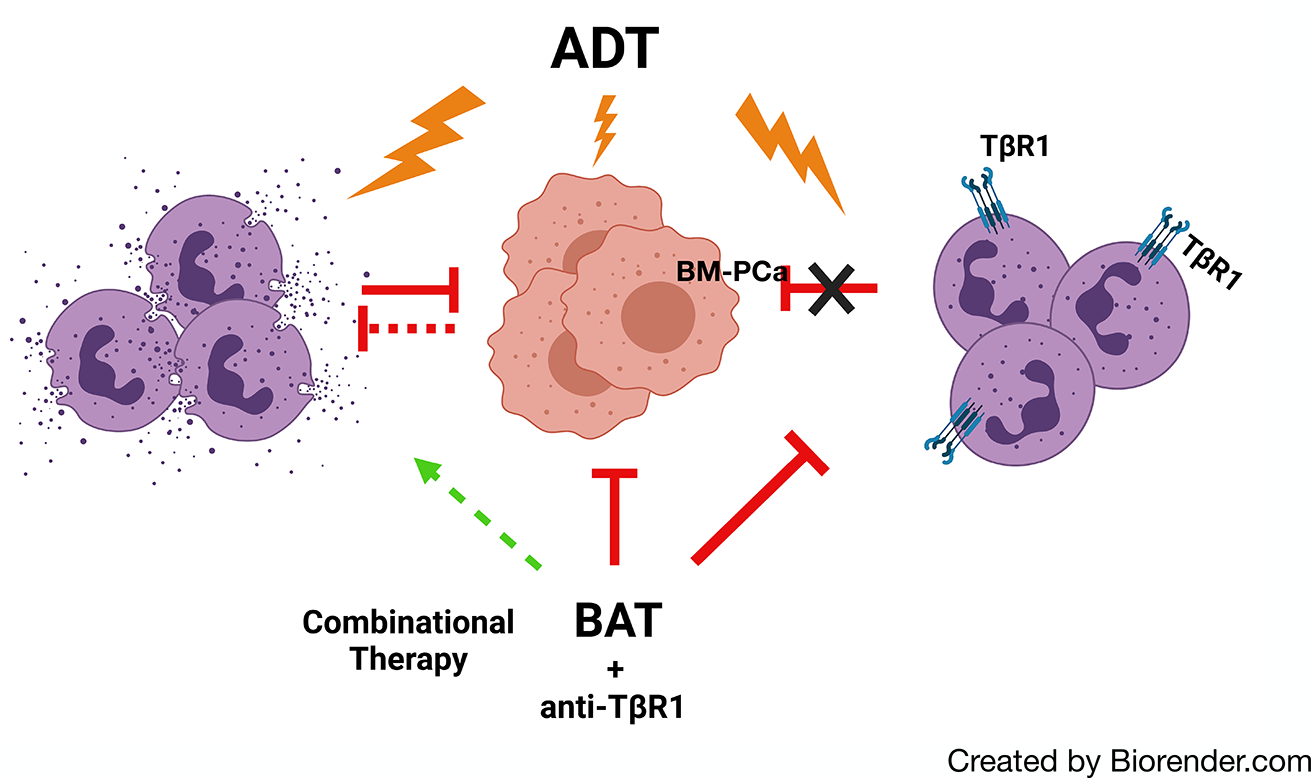
Schematic of androgen regulation and neutrophils in BM-PCa. Bone metastatic prostate cancer (BM-PCa) cells are in the center and bone marrow neutrophils (purple) are left and right of BM-PCa. Red lines represent inhibition, green dashed arrow represent potential signal promotion. BAT = bipolar androgen therapy

## Discussion

Prostate growth is predominantly regulated by androgen signaling; thus, androgen deprivation remains the primary therapeutic approach for treating prostate cancer. However, patients frequently develop resistance to androgen deprivation therapies e.g., develop castration-resistant prostate cancer and the development of bone metastatic disease. In this study, we examined the impact of prostate cancer disease progression on neutrophils to determine their potential as biomarkers for disease progression and therapeutic response. This is the first study to delineate the functional and molecular profile of PMNs in PCa and we reveal important findings suggesting that classical PMN characterization is not parallel to PMN cytotoxicity which is highly dependent on androgen-focused hormone therapy.

Neutrophils/PMNs have been classically ignored in the conversation about immunotherapy due to their propensity to be short-live terminally differentiated cells, most well characterized in infectious diseases, with emerging findings of cancer-related properties that seem to differ depending on tissue and disease context. In preclinical cancer studies, there has been a collection of methods derived and recently described, for proper characterization of tumor-associated neutrophils. These methods include viability (shown to be increased in tumor microenvironments) mobilization/migration, activation status (ROS, NETs and granule release), cell morphology and surface markers (via FACS), and cell cytotoxicity assays. Our findings were in support of other studies showing that several of these described PMN properties are enhanced as the PCa progresses; specifically, with PCa progression PMNs demonstrated extended viability, were mobile, exhibited a pro-inflammatory molecular shift and appeared to be comprised of less mature cells in favor of more immature (and potentially immunosuppressive) populations.

As mentioned, we also identified disease stage-specific differences in PMN populations that may be useful in downstream studies to understand ideal PMN-focused therapeutic interventions. Similar to our previous data showing that cancer soluble factors increase neutrophil viability, there was a significant increase in the viability of all PCa-derived PMNs, specifically of those with localized disease compared to healthy men which suggests a less mature population since mature PMNs are terminally differentiated and lack mitotic ability^33^. In support of this, there changes in maturation markers, such as CD10 which was significantly reduced with progression of disease, along with CD66b. Both CD10 and CD66b have been associated with PMN maturity and are highly expressed on fully differentiated PMNs, compared to immature precursors^34,35^. There was a significant increase in PMNs expressing CD88 (C5aR) in mCRPC compared to mCSPC. Complement C5a is a potent chemoattract for leukocyte migration, promotes granule enzyme release and oxidative burst and thus, often is associated with neutrophil activation^36^. This suggests more activated PMNs in mCRPC compared to less aggressive prostate cancer. It is possible that these cells are highly cytotoxic based on our previous studies and this idea was supported by RNA sequencing data which revealed a gene signature associated neutrophil activation and degranulation, based upon expression of granule enzymes, complement receptors, and genes associated with migration and oxidative burst. This could also suggest a pro-inflammatory activated state, identified in PMNs from the most aggressive stage mCRPC which could be detrimental to other tissues. Additionally, there was an increase in Lox-1 (marker for immunosuppressive cells^37^) positive cells in both cohorts of metastatic patients compared to healthy men and those with localized PCa. However, in total fewer than 20% of collected PMNs expressed Lox-1 suggesting that Lox-1 expression alone may not be an appropriate marker for determining PMN function in BM-PCa. Ongoing studies will be needed to determine the relation of these cells with the androgen-regulated TΒRI-positive cell populations.

However, PMN properties acquired throughout progression did *not* directly correlate with PMN cytotoxicity *ex vivo* against PCa cells and we verified this in a preclinical BAT/androgen regulation trial. Progression of PCa disease to CRPC is associated with dysregulated androgen signaling, including, but not limited to: 1) restoration of AR signaling via increased AR expression, therapy-induced AR-activating mutations, or splice variants (e.g., AR-V7), 2) bypass of AR signaling pathways, such as STAT5 or glucocorticoid receptor (GR) signaling), and 3) AR independent signaling that occurs upon loss of AR expression (i.e., Myc activation)^38^. The majority of PCa patients in our study received one round of ADT to inhibit circulating androgen levels (e.g., GnRH agonist/antagonist, LH agonist) and, upon biochemical recurrence, will receive second-generation androgen inhibitors or ARSIs e.g., Abiraterone or Enzalutamide/Darolutamide/Bicalutamide, respectively. Although second generation androgen inhibition and ARSI prove to be temporarily beneficial for treating disease recurrence in CRPC, immune cells also rely on AR for function and our data demonstrates that ADT specifically impacts neutrophil cytotoxic immune response more so than PCa disease stage.

There have been several studies to demonstrate the impact of AR regulation on the immune system. AR is present on all PMNs and PMN precursor cells within the bone. Previous studies revealed that granulopoiesis i.e., the formation of granulocytes in the bone marrow, is regulated by androgen signaling. Specifically, castration of male mice significantly reduced neutrophil number and this was further verified using conditional AR KO mice, where neutrophil numbers were reduced by 90%^39,40^. In patients, AR inhibition reduces PMN numbers however, castration does not always result in neutropenia suggesting that AR specifically is more important for PMN differentiation than granulopoiesis. This was attributed to AR-mediated granulocyte-colony stimulating factor, the predominant cytokine that stimulates PMN differentiation and proliferation^41,42^. Along these lines, in our BAT trial, we found that AR-inhibited TBNs expressed more SCF/cKit ligand, which has been shown to be associated with immature neutrophils^43^; this was in comparison to mice treated with BAT, which showed little to no SCF/cKit expression, suggesting that AR mediation of neutrophil function may relate to suppression of maturation. Although some of our patients received other non-androgen focused therapies, however a longer more comprehensive study would be needed to examine the importance of therapies specifically for BM-PCa.

Based on known roles for TGFβ in the-bone microenvironment^44–46^ and in regulating TAN phenotypes, we examined the role of TGFβ receptor in androgen-mediated neutrophil response. *In vitro* preclinical data revealed that androgen mediation of neutrophils is predominantly regulated by neutrophil TβRI expression. Canonical TGFβ signaling is propagated through ligand binding of the TβR2 receptor which then forms a heterodimer complex with the kinase receptor, TβRI. TβRI activation results in phosphorylation of Smad2/3 docked at the receptor, dimerization with cytoplasmic Smad4 and translocation into the nucleus for transcriptional regulation through Smad4 binding of Smad-binding elements (SBEs)^47^. Several studies have identified a role for TβRI in cancer immunity, namely as an immunosuppressant and a known regulator of inflammation^48–50^. Recently, Paller et al. showed that TβRI inhibition using the small molecule kinase inhibitor, Galunisertib, combined with a dominant negative TβRI suppressed primary prostate tumor growth and enhanced response to Enzalutamide in the neuroendocrine transgenic adenoma mouse prostate (TRAMP) model^51^. In a separate study, PCa sequencing study, scRNAseq of prostate primary tumors from patients with advanced disease showed an increase in TβRI expression and correlation with enzalutamide resistance demonstrating a connection between AR inhibition and TGFβ signaling in both the epithelial cancer cells as well as stromal cells^52^. It is unclear how androgens regulate TGFβ signaling. A previous study showed that AR regulates TGFβ-mediated signaling through direct association with Smad3, thereby suppressing Smad-mediated transcription. Additionally, we and others have shown TGFβ receptor expression to be regulated by TGFβ signaling^45,46^. These data suggest that ARSIs disrupt the AR-Smad binding which indirectly increases TΒRI expression.

Although TGFβ is ubiquitously expressed, mouse scRNAseq data revealed expression predominantly in granulocytic cells in the bone marrow. Further, TβRI appears to be expressed more abundantly in neutrophils in patient bone metastases, compared to matched liver metastases suggesting that systemic TβRI inhibition may be feasible for enhancing neutrophil anti-tumor response in bone. In support of this, we show that combined systemic AR and TβRI inhibition suppressed tumor growth of AR-independent prostate cancer in bone. Further, we have found that AR-negative PC3 cells are resistant to neutrophil killing; however, neutrophil-specific loss of TβRI expression was able to sensitize PC3 to neutrophil killing and enzalutamide treatment suggesting that both neutrophil TβRI suppression combined with AR inhibition, significantly enhances neutrophil cytotoxicity to even the resistant BM-PCa. Further studies will be required to determine the mechanisms of TβRI and AR-mediated neutrophil function in the tumor-bone microenvironment.

Collectively, these data demonstrate that peripheral blood PMNs can give insight into PCa disease progression, particularly in the context of bone-metastatic disease. Importantly, we also identified reveal a novel finding that neutrophil anti-tumor immune response is suppressed by ADT through TβRI (Figure 7). With the benefit of current standard-of-care ADT on PCa progression, this information can be used as a two-pronged approach, leveraging TβRI as a viable target for enhancing neutrophil anti-tumor response in the presence of ADT.

## Supporting information

Alsamraae et al. Supplemental Data

## Acknowledgements

The authors thank Dr. Yusuke Shiozawa for kindly providing the RM1 cells and Dr. Mathias Gunzer for providing the Catchup mice in this manuscript. LMC was supported by a Research Scholar Grant (RSG-19-127-01-CSM) from the American Cancer Society. This work was supported in part by the Flow Cytometry, Small Animal Imaging Laboratory, and Molecular Biology cores at UNMC. The authors acknowledge the UNMC Multiphoton Intravital & Tissue Imaging (MITI) Core (RRID:SCR_022478) supported by P30GM127200, P20GM130447, P30CA036727, and the Nebraska Research Initiative. The UNMC Genomics Core Facility receives partial support from the National Institute for General Medical Science (NIGMS) INBRE - P20GM103427-19, as well as the National Cancer Institute. The UNMC Flow Cytometry Research Facility is administrated through the Office of the Vice Chancellor for Research and supported by state funds from the Nebraska Research Initiative (NRI) and The Fred and Pamela Buffett Cancer Center’s National Cancer Institute Cancer Support Grant. Major instrumentation has been provided by the Office of the Vice Chancellor for Research, The University of Nebraska Foundation, the Nebraska Banker’s Fund, and by the NIH-NCRR Shared Instrument Program. Tissue acquisition in the University of Washington Prostate Cancer Donor Rapid Autopsy Program was supported by the Pacific Northwest Prostate Cancer SPORE (P50CA97186). This research was supported, in part, by the Intramural Research Program of the NIH, National Cancer Institute, Center for Cancer Research (CCR) and Cancer Innovation Laboratory (CIL).

