## Supplementary material for "Androgen-mediated TGFβ expression suppresses anti-tumor neutrophil response in bone metastatic prostate cancer": Alsamraae et al. Supplemental Data

### Supplemental Figures

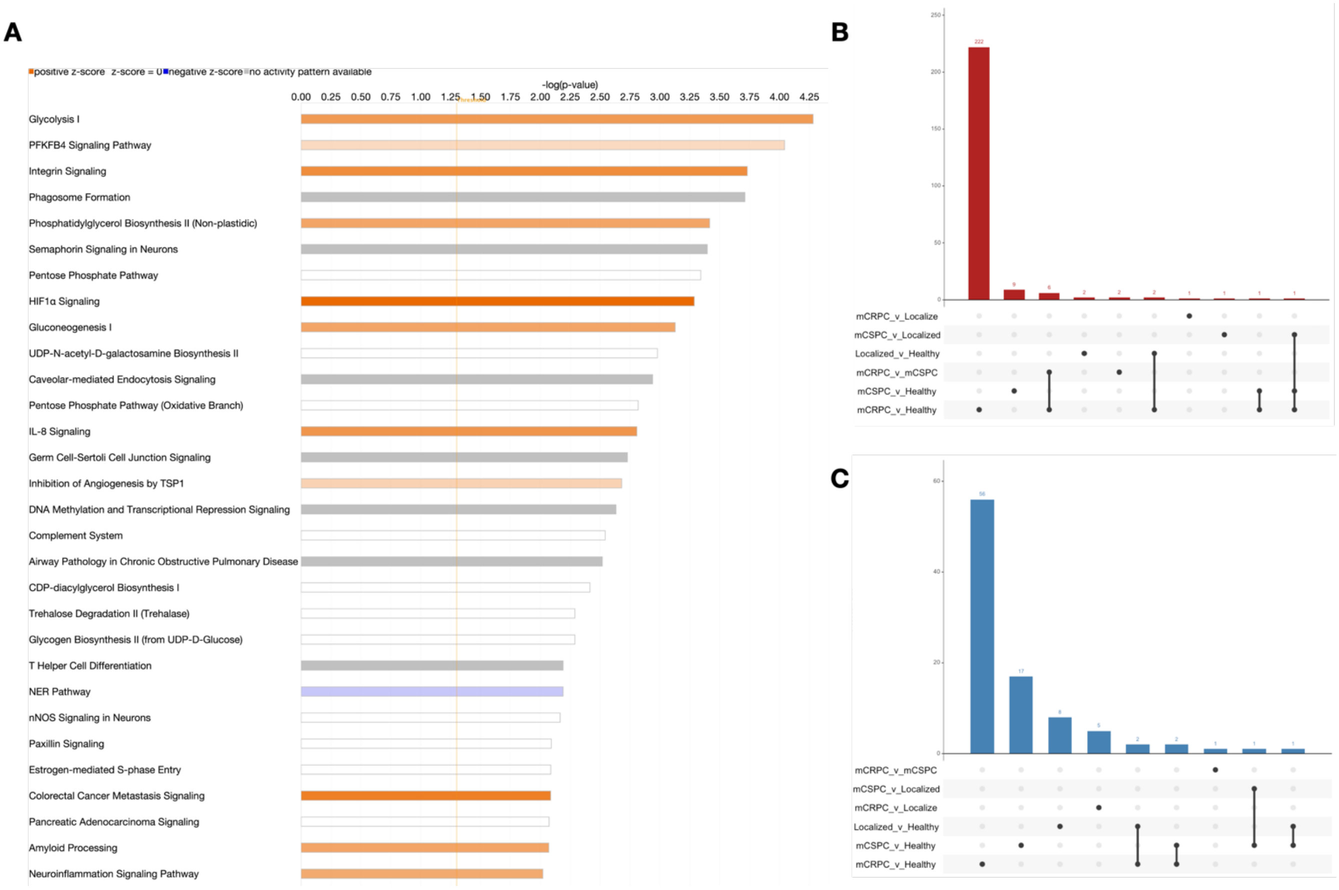

**Supplemental Figure 1. Analysis of bulk RNA sequencing of PMNs in PCa progression.** (A) Ingenuity Pathway Analysis (IPA) canonical pathway activation in mCRPC compared to healthy PMNs. (B) Upset plot of up-regulated PMN genes in PCa. (C) Upset plot of down-regulated PMN genes in PCa.

**A**

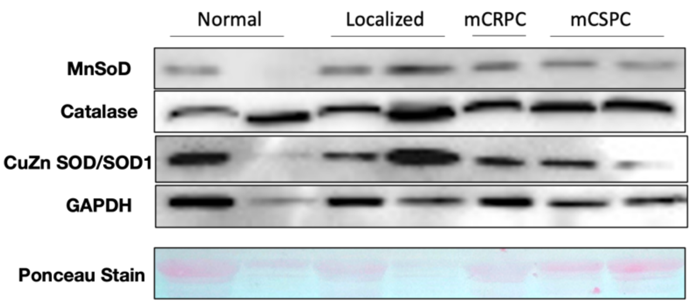

**B**

| Group | patient # | SOD1 | | SOD | | Catalase |
|---|---|---|---|---|---|---|
| Normal | 3203974 | yes | | yes | | yes |
| Normal | 2512220 | yes | | yes | | yes |
| Normal | 334725 | no | | yes | | yes |
| Normal | 2491212 | no | | yes | | yes |
| Normal | 397634 | yes | | yes | | yes |
| Normal | 2424678 | yes | | yes | | yes |
| Normal | 453049 | yes | | yes | | yes |
| Normal | 2045394 | yes | | yes | | yes |
| Localized | 3122723 | yes | | yes | | yes |
| Localized | 2041622 | yes | | yes | | yes |
| Localized | 15354 | no | | yes | | yes |
| Localized | 2485834 | no | | yes | | yes |
| Localized | 1258461 | yes | | yes | | yes |
| Localized | 3158761 | yes | | yes | | yes |
| Localized | 3670961 | yes | | yes | | yes |
| Localized | 1153600 | yes | | yes | | yes |
| mCRPC | 490334 | yes | | yes | | yes |
| mCRPC | 214884 | no | | yes | | yes |
| mCRPC | 2478651 | no | | yes | | yes |
| mCRPC | 3759009 | no | | yes | | yes |
| mCRPC | 2263201 | yes | | yes | | yes |
| mCRPC | 2337963 | no | | yes | | yes |
| mCRPC | 6259 | yes | | yes | | yes |
| mCSPC | 2043564 | no | | yes | | yes |
| mCSPC | 1153640 | no | | yes | | yes |
| mCSPC | 1404067 | no | | yes | | yes |
| mCSPC | 1102755 | no | | yes | | yes |
| mCSPC | 77348 | yes | | yes | | yes |
| mCSPC | 294350 | no | | yes | | yes |
| mCSPC | 74390 | yes | | yes | | yes |
| mCSPC | 2234624 | yes | | yes | | yes |

**Supplemental Figure 2. Protein expression of PMN antioxidants.** (A) Representative western blot of PMN antioxidants. GAPDH was loaded as a control. (B) Table of represented antioxidants per PCa disease stage; yes/no for positive or negative detection per patient sample (n=8/group).

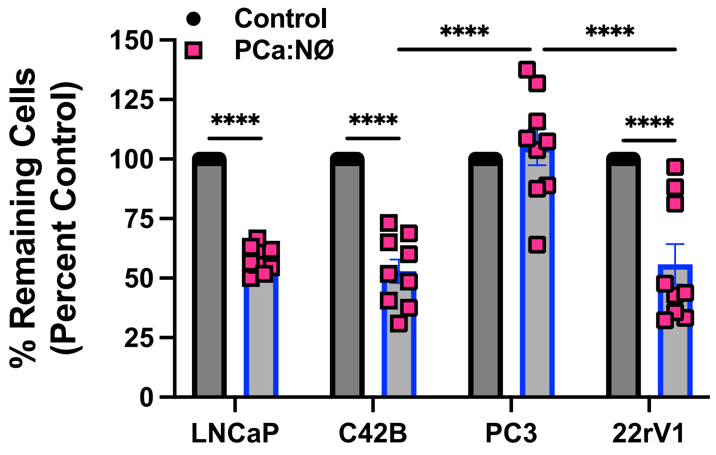

**Supplemental Figure 3. Patient PMN co-culture of BM-PCa.** Graph shows patient PMN co-culture with LNCaP, C42B, PC3 and 22rV1 cells (n=3 patients).

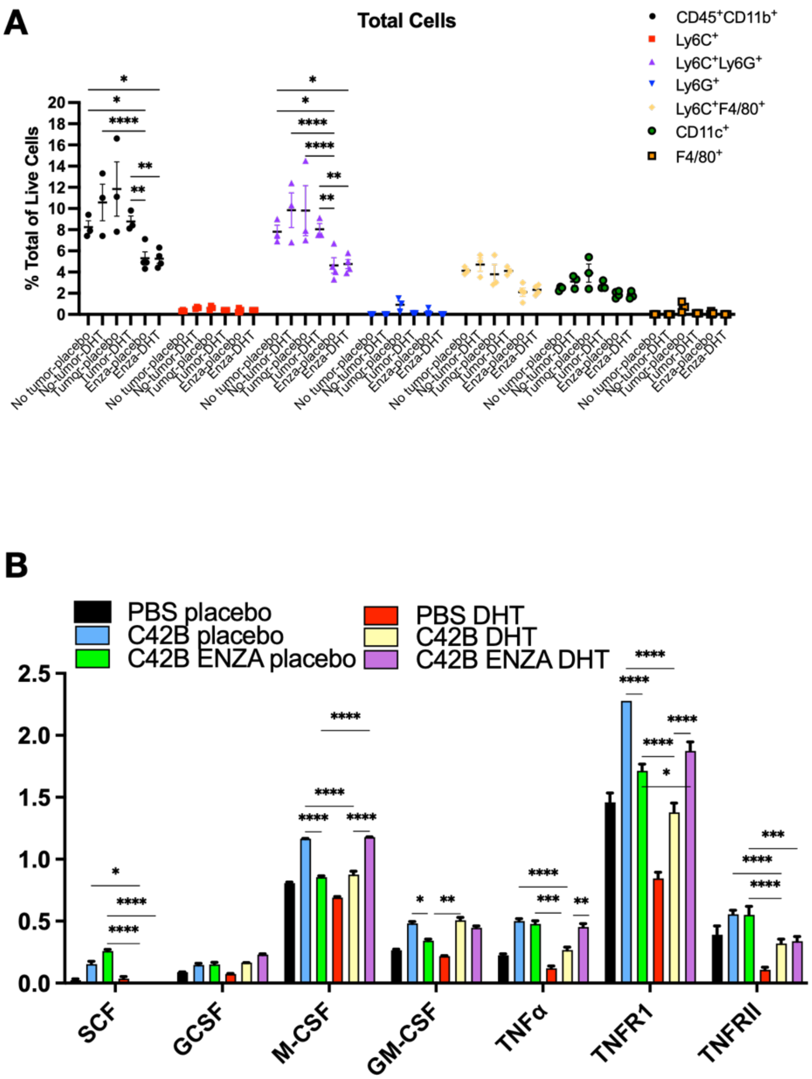

**Supplemental Figure 4. Flow cytometry and cytokine array of mouse bone marrow and TBNs in response to androgen regulation.** (A) Flow cytometry of mouse bone marrow from ADT and BAT preclinical trial. Graph shows percent positive for respective myeloid markers per live cells. (B) Protein was isolated from mouse TBNs from treated mice and cytokines probed via membrane array.

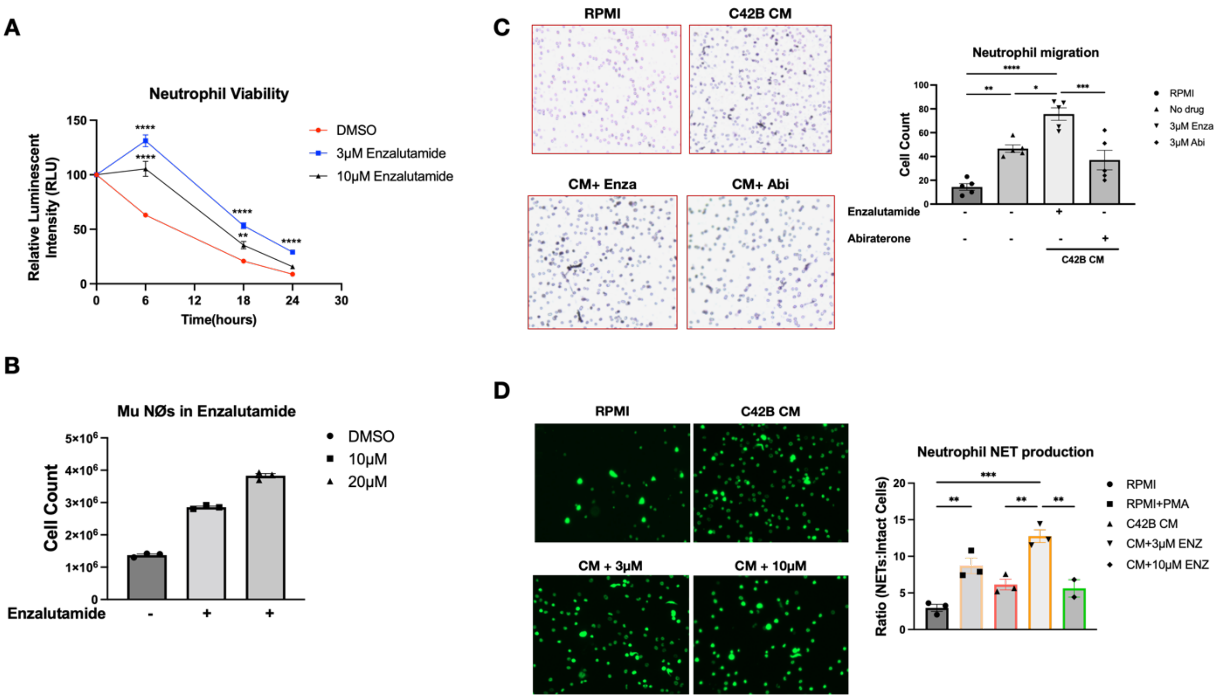

**Supplemental Figure 5. Androgen regulation of bone marrow neutrophil functions.** Mouse bone marrow neutrophils were treated with enzalutamide or abiraterone and neutrophil properties measured. (A) Enzalutamide was added to the media and neutrophil viability using CellGro assay. (B) Neutrophil cell counts after overnight treatment with enzalutamide. (C) Boyden chamber migration assay performed one hour after 30-minute ARSI treatment. (D) Sytox Green NETosis assay of neutrophils after 30-minute treatment with ARSIs.

### Supp. Figure 6

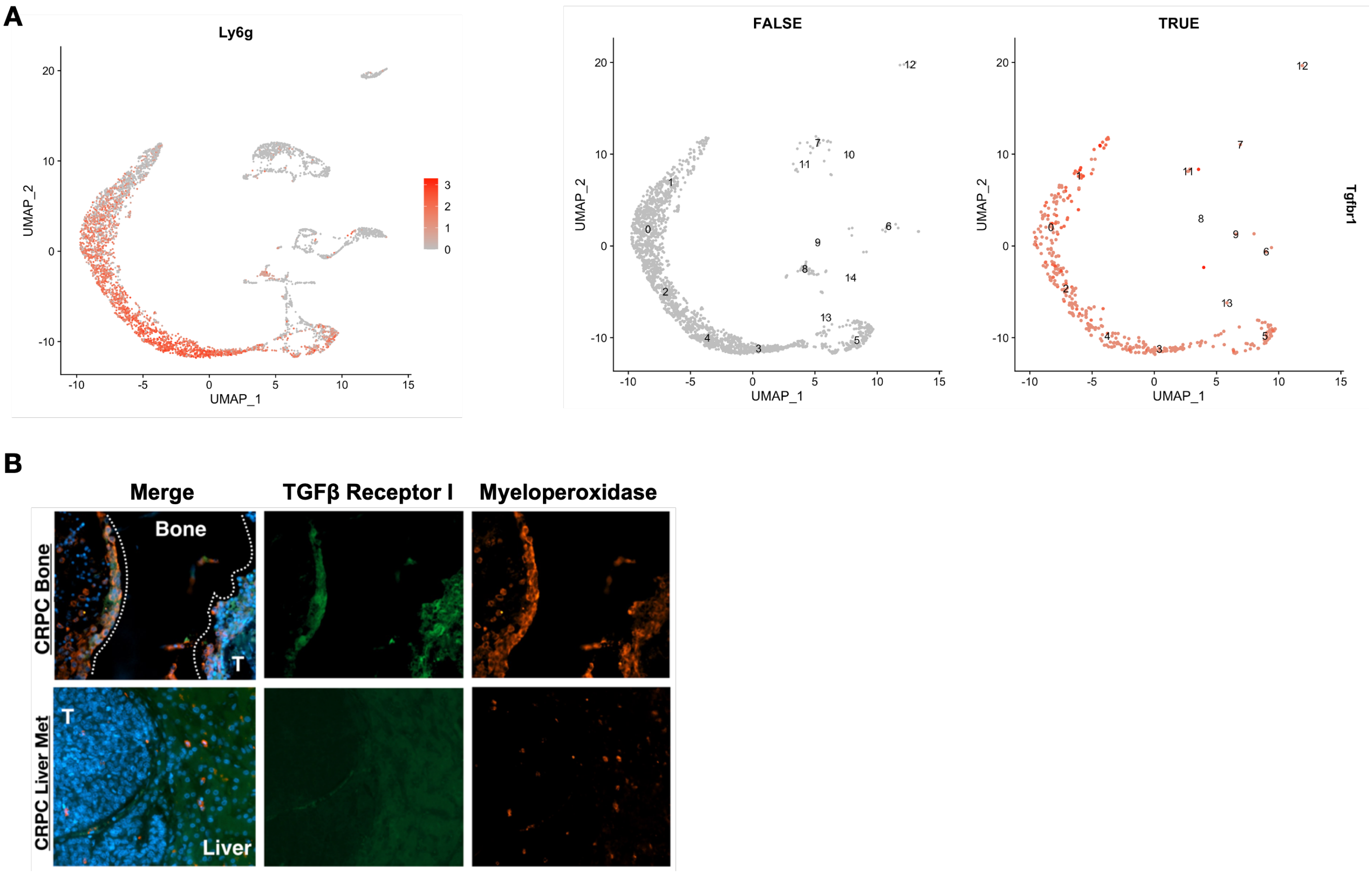

**Supplemental Figure 6. TβRI expression in mouse bone marrow and patient bone biopsies.** (A) Left, UMAP of Ly6G$^+$ bone marrow cells. Right, UMAP of TβRI-positivity; false = Ly6G$^+$, TβRI$^-$ cells, positive = Ly6G$^+$, TβRI$^+$ cells. (B) Representative immunofluorescence images (20x) of mPCa patient matched bone (top) and liver (bottom) metastases. Neutrophils are identified by myeloperoxidase (orange) and TβRI (green) expression analyzed. Nuclei were stained with DAPI (blue).

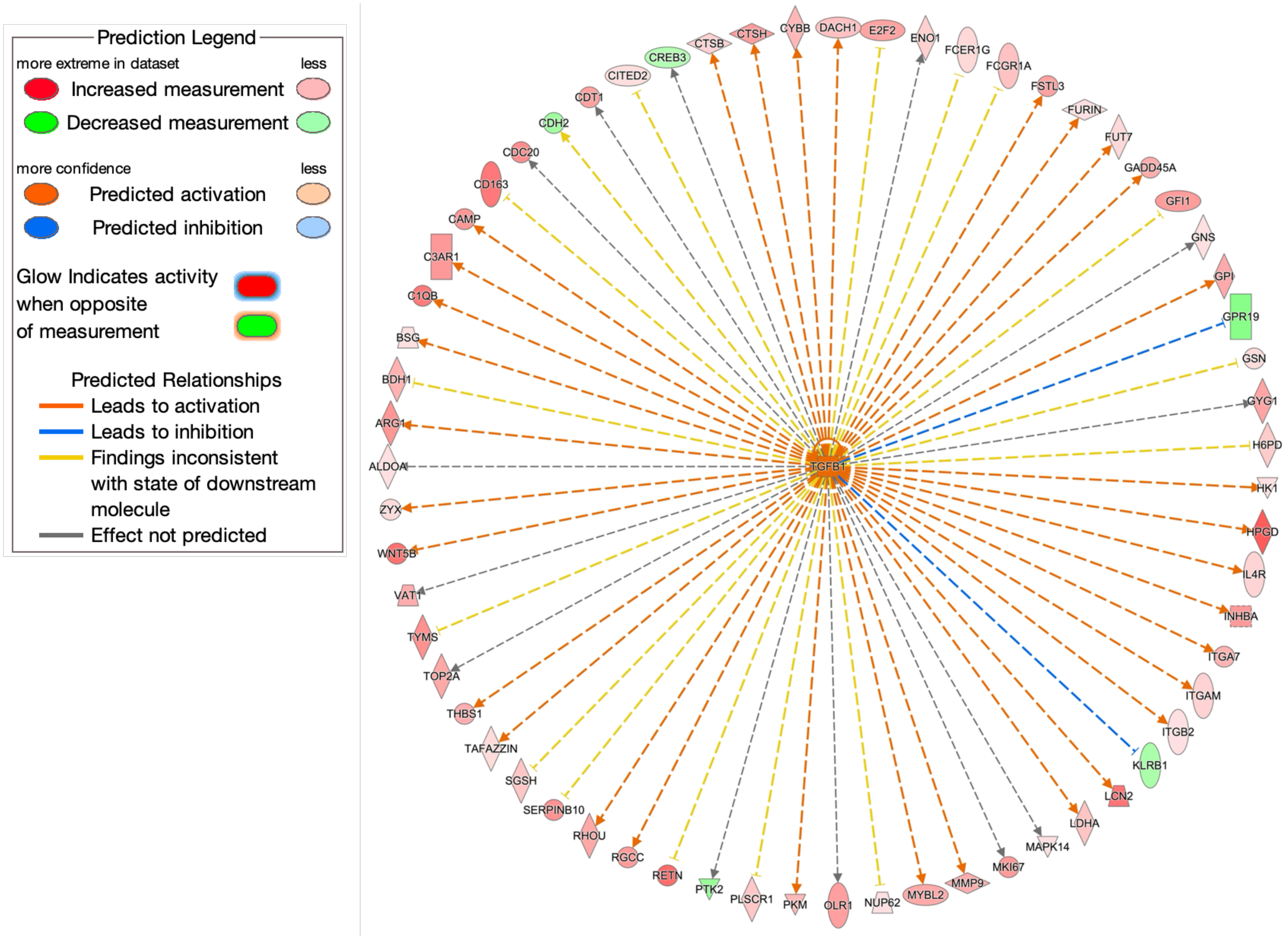

**Supplemental Figure 7. TGFβ in Patient PMNs.** TGFβ was identified as an upstream mediator of gene regulation in mCRPC PMNs compared to healthy patients, using IPA software. Image shows mCRPC PMN downstream targets of TGFβ and type (up/down) described in legend.

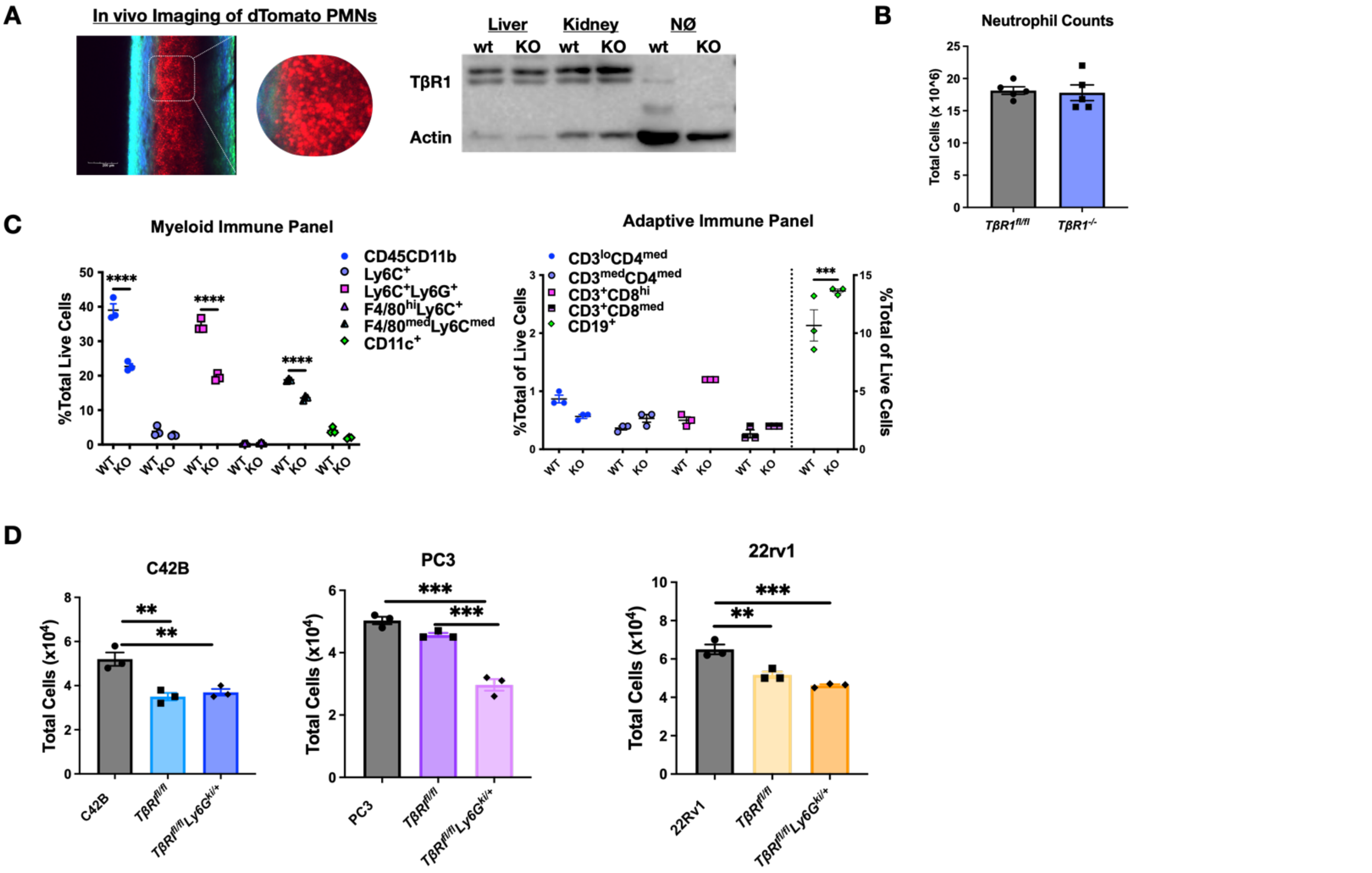

**Supplemental Figure 8. TβRI knockout mouse characteristics.** (A) Left, two-photon microscopy of bone marrow of Catchup mouse model. Right, western blot of TβRI in tissues from Catchup/ *TβRI* $^{flox/flox}$ crossed mice. (B) Neutrophil counts post-isolation from mouse bone marrow. (C) Flow cytometry of myeloid cell markers (left) and adaptive immune markers (right) from mouse bone marrow of wildtype TβRI and TβRI-null mice. (D) Co-culture of mouse neutrophils with PCa cell lines.

**Supplemental Table 1. Flow Cytometry Antibodies**

| **Human** | **RRID** |
|---|---|
| APC/Cyanine7 anti-mouse/human CD11b | RRID:AB_830641 |
| PE/Cyanine7 anti-human CD14 | RRID:AB_389352 |
| FITC anti-human CD15 (SSEA-1) | RRID:AB_314195 |
| APC anti-human CD16 | RRID:AB_314211 |
| PerCP/Cyanine5.5 anti-human CD10 | RRID:AB_10643591 |
| PerCP/Cyanine5.5 anti-human CD66b | RRID:AB_2077856 |
| PE anti-human CD88 (C5aR) | RRID:AB_1877226 |
| APC anti-human HLA-DR | RRID:AB_314688 |
| PE anti-human LOX-1 | RRID:AB_2562180 |
| **Mouse** | |
| *Myeloid/Innate Panel* | |
| APC/Cyanine7 anti-mouse CD45 | RRID:AB_312980 |
| FITC anti-mouse/human CD11b | RRID:AB_312788 |
| PE anti-mouse Ly6G | RRID:AB_1186104 |
| PerCP/Cyanine5.5 anti-mouse Ly6C | RRID:AB_1659242 |
| PE/Cyanine7 anti-mouse F4/80 | RRID:AB_893490 |
| APC anti-mouse CD11c | RRID:AB_313778 |
| *Adaptive Panel* | |
| APC anti-mouse CD3 | RRID:AB_2561455 |
| FITC anti-mouse CD4 | RRID:AB_312690 |
| PE/Cyanine7 anti-mouse CD8a | RRID:AB_312760 |
| APC/Cyanine7 anti-mouse CD19 | RRID:AB_830706 |

**Supplemental Table 2. PCR Primer Sequences**

| **Mouse Primer Sequences** | |
|---|---|
| muTβRI | FWD 5’-CGCTCTGTCCACGGCAAG-3’<br>REV 5’-TCATGTCTCACAGCAAGTCCC-3’ |
| muTβRII | FWD 5’-GGCCAAGCTGAAGCAGAA-3’<br>REV 5’-GGATGTTCTCGTGTTTCAGGTT-3’ |
| TNFalpha | FWD 5’- TAGCCCACGTCGTAGCAAAC-3’<br>REV 5’- GCAGCCTTGTCCCTTGAAGA-3’ |
| **Human Primer Sequences** | |
| TβRI | FWD 5’-CCCAGCATCTGCAAAGCTC-3’<br>REV 5’-GTCAATGTACAGCTGCCGCA-3’ |
| TβR2 | FWD 5’-ACTTTATTCTGGAAGCTGCT-3’<br>REV 5’-GCTGATGCCTGTCACTTGAA-3' |
| Human/Mouse AR | FWD 5’- CATGTGGAAGCTGCAAGGTCT-3’<br>REV 5’- TCTGTTTCCCTTCAGCGGC-3’ |
